# Omicron breakthrough infection drives cross-variant neutralization and memory B cell formation

**DOI:** 10.1101/2022.04.01.486695

**Authors:** Jasmin Quandt, Alexander Muik, Nadine Salisch, Bonny Gaby Lui, Sebastian Lutz, Kimberly Krüger, Ann-Kathrin Wallisch, Petra Adams-Quack, Maren Bacher, Andrew Finlayson, Orkun Ozhelvaci, Isabel Vogler, Katharina Grikscheit, Sebastian Hoehl, Udo Goetsch, Sandra Ciesek, Özlem Türeci, Ugur Sahin

## Abstract

Omicron is the evolutionarily most distinct SARS-CoV-2 variant (VOC) to date and displays multiple amino acid alterations located in neutralizing antibody sites of the spike (S) protein. We report here that Omicron breakthrough infection in BNT162b2 vaccinated individuals results in strong neutralizing activity not only against Omicron, but also broadly against previous SARS-CoV-2 VOCs and against SARS-CoV-1. We found that Omicron breakthrough infection mediates a robust B cell recall response, and primarily expands preformed memory B cells that recognize epitopes shared broadly by different variants, rather than inducing new B cells against strictly Omicron-specific epitopes. Our data suggest that, despite imprinting of the immune response by previous vaccination, the preformed B cell memory pool has sufficient plasticity for being refocused and quantitatively remodeled by exposure to heterologous S protein, thus allowing effective neutralization of variants that evade a previously established neutralizing antibody response.

**One Sentence Summary:** Breakthrough infection in individuals double- and triple-vaccinated with BNT162b2 drives cross-variant neutralization and memory B cell formation.

## Introduction

Containment of the current COVID-19 pandemic requires the generation of durable and sufficiently broad immunity that provides protection against circulating and future variants of SARS-CoV-2. The titer of neutralizing antibodies to SARS-CoV-2, and the binding of antibodies to the spike (S) glycoprotein and its receptor-binding domain (RBD) are considered correlates of protection against infection (*1, 2*). Currently available vaccines are based on the ancestral Wuhan-Hu-1 strain and induce antibodies with a neutralizing capacity that exceeds the breadth elicited by infection with the Wuhan strain, or with variants of concern (VOCs) (*3*). However, protective titers wane over time (*4–7*) and routine booster vaccinations are thought to be needed to trigger recall immunity and maintain efficacy against new VOCs (*8–11*).

Long-lived memory B (B_MEM_) cells are the basis for the recall response upon antigen reencounter either by infection or booster vaccination. They play an important role in the maintenance and evolution of the antiviral antibody response against variants, since low-affinity selection mechanisms during the germinal center reaction and continued hypermutation of B_MEM_ cells expand the breadth of viral variant recognition over time (*12, 13*).

How vaccine-mediated protective immunity will evolve over time and will be modified by iterations of exposure to COVID-19 vaccines and infections with increasingly divergent viral variants, remains poorly understood and is of particular relevance with the emergence of antigenically distinct VOCs. Omicron is the evolutionary most distant reported VOC with a hitherto unprecedented number of amino acid alterations in its S glycoprotein, including at least 15 amino acid changes in the RBD and extensive changes in the N-terminal domain (NTD) (*14*). These alterations are predicted to affect most neutralizing antibody epitopes (*15–18*). In addition, Omicron is highly transmissible, and its sublineages BA.1 and BA.2 have spread rapidly across the globe, outcompeting Delta within weeks to become the dominant circulating VOC (*19, 20*).

To date, over 1 billion people worldwide have been vaccinated with the mRNA-based COVID-19 vaccine BNT162b2 and have received the primary 2-dose series or further boosters (*21*). This vaccine is contributing substantially to the pattern of population immunity in many regions on which further immune editing and effects of currently spreading variants will build upon.

To characterize the effect of Omicron breakthrough infection on the magnitude and breadth of serum neutralizing activity and B_MEM_ cells, we studied blood samples from individuals that were double- or triple-vaccinated with BNT162b2.

As an understanding of the antigen-specific B cell memory pool is a critical determinant of an individual’s ability to respond to newly emerging variants, our data will help to guide future vaccine development.

## Results and discussion

### Cohorts and sampling

Blood samples have been sourced from the biosample collection of BNT162b2 vaccine trials, and a biobank of prospectively collected samples from vaccinated individuals with subsequent SARS-CoV-2 Omicron breakthrough infection. Samples were selected to investigate biomarkers in four independent groups, namely individuals who were (i) double- or (ii) triple-vaccinated with BNT162b2 without a prior or breakthrough infection at the time of sample collection (BNT162b2^2^, BNT162b2^3^) and individuals who were (iii) double- or (iv) triple-vaccinated with BNT162b2 and who experienced breakthrough infection with the SARS-CoV-2 Omicron variant after a median of approximately 5 months or 4 weeks, respectively (BNT162b2^2^ + Omi, BNT162b2^3^ + Omi) (see materials and methods). Immune sera were used to characterize Omicron infection-associated changes to the magnitude and the breadth of serum neutralizing activity. PBMCs were used to characterize the VOC-specificity of peripheral B_MEM_ cells recognizing the respective full-length SARS-CoV-2 S protein or its RBD (Fig. 1, Tables S1 to S3).

**Fig. 1.**
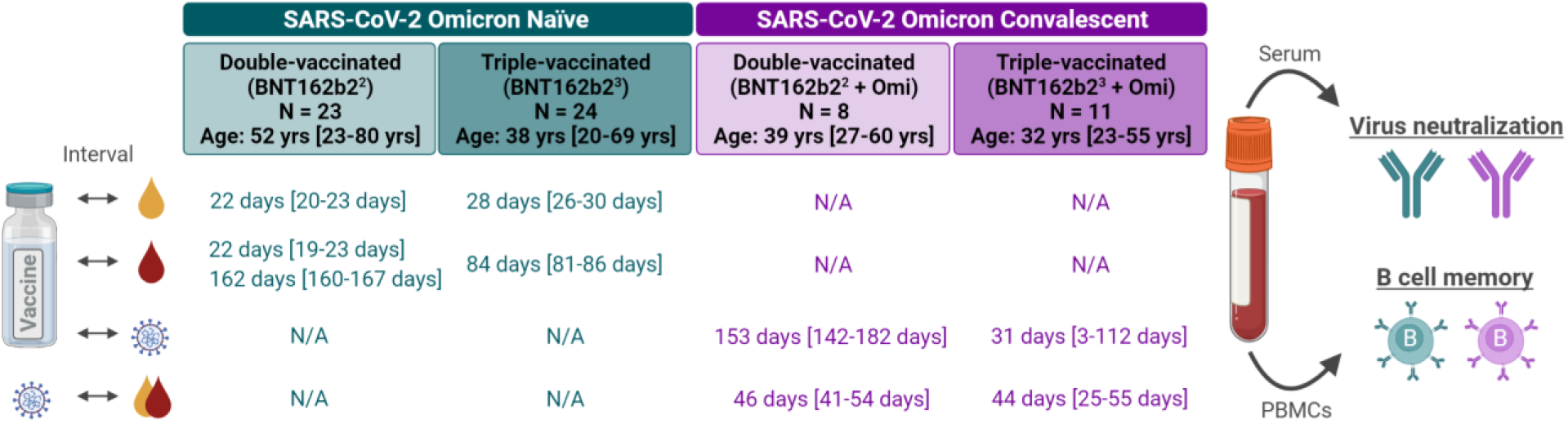
Cohorts, sampling and experimental setup. Blood samples were drawn from four cohorts: Omicron-naïve individuals double- or triple-vaccinated with BNT162b2 (light and dark green), and individuals double- or triple-vaccinated with BNT162b2 that subsequently had a breakthrough infection with Omicron (light and dark purple). PBMCs (red) and sera (yellow) were isolated in the Omicron-naïve cohorts at the time-points indicated following their most recent vaccination; for convalescent cohorts, the time from their most recent vaccination to Omicron infection, and infection to PBMC and serum isolation are indicated (all values specified as median-range). Serum neutralizing capacity was assessed using a pseudovirus and live virus neutralization test; SARS-CoV-2 spike-specific B_MEM_ cells were assessed via a flow cytometry-based B cell phenotyping assay using bulk PBMCs. N/A, not applicable; Schematic was created with BioRender.com

### Omicron breakthrough infection of BNT162b2 double- and triple-vaccinated individuals induces broad neutralization of Omicron BA.1, BA.2 and other VOCs

To evaluate the neutralizing activity of immune sera, we used two orthogonal test systems: a well-characterized pseudovirus neutralization test (pVNT) (*22, 23*) to investigate the breadth of inhibition of virus entry in a propagation-deficient set-up, as well as a live SARS-CoV-2 neutralization test (VNT) designed to evaluate neutralization during multicycle replication of authentic virus with the antibodies maintained throughout the entire test period. For the former, we applied pseudoviruses bearing the S proteins of Omicron sublineages BA.1 or BA.2, other SARS-CoV-2 VOCs (Wuhan, Alpha, Beta, Delta) to assess breadth and of SARS-CoV (herein referred as SARS-Cov-1) to detect potential pan-Sarbecovirus neutralizing activity (*24*).

As reported previously (*22, 25, 26*), in Omicron-naïve double-vaccinated individuals 50% pseudovirus neutralization (pVN_50_) geometric mean titers (GMTs) of Beta and Delta VOCs were reduced, and neutralization of both Omicron sublineages was virtually undetectable (Fig. 2a, fig. S1a, Table S4). In Omicron-naïve triple-vaccinated individuals, pVN_50_ GMTs against all tested VOCs were substantially higher with robust neutralization of Alpha, Beta and Delta. While GMTs against Omicron BA.1 were significantly lower compared to Wuhan (GMT 160 vs 398), titers against Omicron BA.2 were also considerably reduced at 211 (Fig. 2a, fig. S1a, Table S5). Thus, triple vaccination induced a similar level of neutralization against the two Omicron sublineages.

**Fig. 2.**
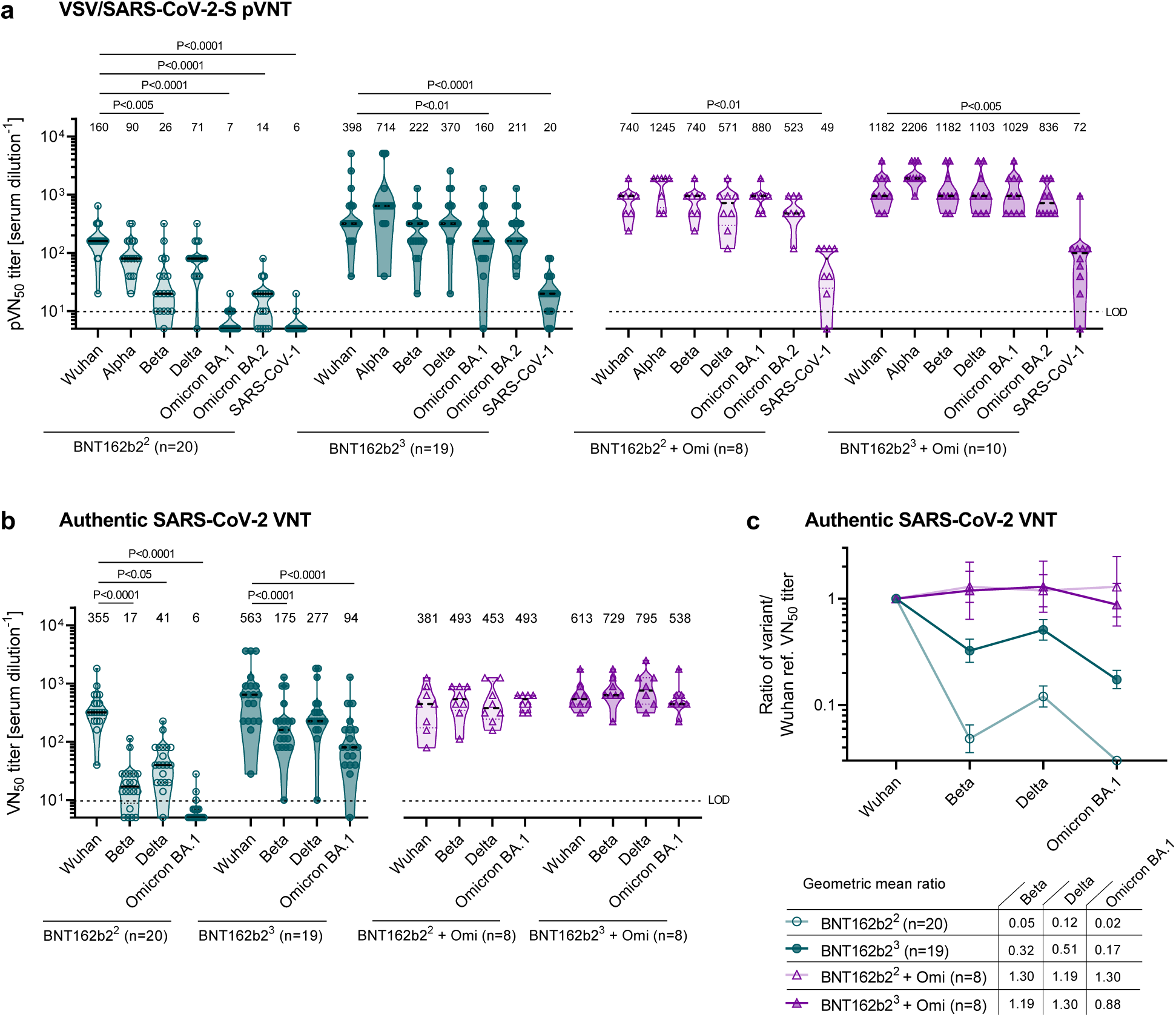
Omicron breakthrough infection of BNT162b2 double- and triple-vaccinated individuals induces broad neutralization of Omicron BA.1, BA.2 and other VOCs. Serum was drawn from double-vaccinated individuals (BNT162b2^2^) at 22 days after the second dose (green, open circles), from triple-vaccinated individuals (BNT162b2^3^) at 28 days after the third dose (green, closed circles), from double-vaccinated individuals with Omicron breakthrough infection (BNT162b2^2^ + Omi) at 46 days post-infection (purple, open triangles), and from triple-vaccinated individuals and Omicron breakthrough infection (BNT162b2^3^ + Omi) at 44 days post-infection (purple, closed triangles). Serum was tested in duplicate; 50% pseudovirus neutralization (pVN_50_) geometric mean titers (GMTs) (in a), 50% virus neutralization (VN_50_) GMTs (in b), and the geometric mean ratio of SARS-CoV-2 variant of concern (VOC) and Wuhan VN_50_ GMTs (in c) were plotted. For titer values below the limit of detection (LOD), LOD/2 values were plotted. Values above violin plots represent group GMTs. The non-parametric Friedman test with Dunn’s multiple comparisons correction was used to compare Wuhan neutralizing group GMTs with titers against the indicated variants and SARS-CoV-1. Multiplicity-adjusted p values are shown. (**a**) pVN_50_ GMTs against Wuhan, VOC and SARS-CoV-1 pseudovirus. (**b**) VN_50_ GMTs against live SARS-CoV-2 Wuhan, Beta, Delta and Omicron BA.1. (**c**) Group geometric mean ratios with 95% confidence intervals for all cohorts shown in b. PBMC samples from double-vaccinated individuals (BNT162b2^2^) at 22 days after the second dose (green, open squares) and 5 months after the second dose (green, open circles), from triple-vaccinated individuals (BNT162b2^3^) at 84 days after the third dose (green, closed circles), from double-vaccinated individuals with Omicron breakthrough infection (BNT162b2^2^ + Omi) at 46 days post-infection (purple, open triangles), and from triple-vaccinated individuals with Omicron breakthrough infection (BNT162b2^3^ + Omi) at 44 days post-infection (purple, closed triangles) were analyzed via flow cytometry for SARS-CoV-2-specific B_MEM_ cell (B_MEM_ – CD3-CD19+CD20+IgD-CD38^int/low^) frequencies via B cell bait staining. (**a**) Schematic of one-dimensional staining of B_MEM_ cells with fluorochrome-labeled SARS-CoV-2 S protein tetramer bait allowing discrimination of variant recognition. Frequencies of Wuhan or VOC full-length S protein-(**b**) and RBD-(**c**) specific B_MEM_ cells for the four groups of individuals were analyzed. Variant-specific B_MEM_ cell frequencies were normalized to Wuhan frequencies for S protein (**d**) and RBD-(**e**) binding. (**f**) Depicts the frequency ratios of RBD protein specific B_MEM_ cells over full-length S protein-specific B_MEM_ cells. Schematic in (a) was created with BioRender.com

Omicron breakthrough infection had a marked effect on magnitude and breadth of the neutralizing antibody response of both double- and triple-vaccinated individuals, with slightly higher pVN_50_ GMTs observed in the triple-vaccinated individuals (Fig. 2a, fig. S1b, Table S6). The pVN_50_ GMT of double-vaccinated individuals with breakthrough infection against Omicron BA.1 and BA.2 was more than 100-fold and 35-fold above the GMTs of Omicron-naïve double-vaccinated individuals. Immune sera from double-vaccinated individuals with breakthrough infection had broad neutralizing activity, with higher pVN_50_ GMTs against Beta and Delta than observed in Omicron-naïve triple-vaccinated individuals (GMT 740 vs. 222 and 571 vs. 370).

The effect of Omicron breakthrough infection on the neutralization of Omicron BA.1 and BA.2 pseudovirus was less pronounced when looking at triple-vaccinated individuals (approximately 7-fold and 4-fold increased neutralization compared to Omicron-naïve triple-vaccinated individuals). pVN_50_ GMTs against Omicron BA.1, BA.2 and Delta were 1029, 836 and 1103 in triple-vaccinated Omicron breakthrough individuals as compared to 160, 211 and 370 in the Omicron-naïve triple-vaccinated. GMTs against all SARS-CoV-2 VOCs, including Beta and Omicron, were close to titers against the Wuhan reference, while noticeably reduced in triple-vaccinated Omicron-naïve individuals.

Likewise, while sera from vaccinated Omicron-naïve individuals had no detectable or only poor pVN_50_ titers against the phylogenetically more distant SARS-CoV-1, convalescent sera of double- and even more markedly of triple-vaccinated Omicron infected individuals robustly neutralized SARS-CoV-1 pseudovirus (Fig. 2a, Table S4 to S6). Nine out of 18 breakthrough infected individuals (four double-vaccinated and five triple-vaccinated) had SARS-CoV-1 pVN_50_ GMTs comparable to or above those against the Wuhan reference in Omicron-naïve double-vaccinated individuals (GMT≥120, Table S4 and S6).

Authentic live SARS-CoV-2 virus neutralization assays conducted with Wuhan, Beta, Delta and Omicron BA.1 confirmed these observations (Fig 2b, fig. S1c, d, Tables S7 to S9). In BNT162b2 double- and triple-vaccinated individuals, Omicron infection was associated with a strongly increased neutralizing activity against Omicron BA.1 with 50% virus neutralization (VN_50_) GMTs in the same range as against the Wuhan strain (Fig 2b; GMT 493 vs. 381 and GMT 538 vs. 613). Similarly, Omicron convalescent double- and triple-vaccinated individuals showed comparable levels of neutralization against other variants as well (e.g. GMT 493 and 729 against Beta), indicating a wide breadth of neutralizing activity.

In aggregate, these data demonstrate that SARS-CoV-2 Omicron breakthrough infection induces neutralization activity of profound breadth in vaccine-experienced individuals, a finding further supported by the calculated ratios of VN_50_ GMTs against the Wuhan strain and SARS-CoV-2 VOCs (Fig. 2c). While double- and to a lesser extent also triple-BNT162b2 vaccinated Omicron-naïve individuals displayed marked differences in neutralization proficiency against VOCs, neutralization activity of Omicron convalescent subjects was leveled to almost the same range of high performance against all variant strains tested.

Likewise, Omicron breakthrough infection had a similarly broad neutralization augmenting effect in individuals vaccinated with other approved COVID-19 vaccines or heterologous regimens (fig. S2, Table S11).

### B_MEM_ cells of BNT162b2 double- and of triple-vaccinated individuals broadly recognize VOCs and are further boosted by Omicron breakthrough infection

Next, we investigated the phenotype and quantity of SARS-CoV-2 S protein specific B cells in these individuals. To this aim, we employed flow cytometry-based B cell phenotyping assays for differential detection of variant-specific S protein-binding B cells in bulk PBMCs. We found that all S protein- and RBD-specific B cells in the peripheral blood were of a B_MEM_ phenotype (B_MEM_; CD20^high^CD38^int/neg^, fig. S3), as antigen-specific plasmablasts or naïve B cells were not detected (data not shown). The assays therefore allowed us to differentiate for each of the SARS-CoV-2 variants between B_MEM_ cells recognizing the full S protein or its RBD that is a hotspot for amino acid alterations, and variant-specific antigenic epitopes (*15, 16*) (Fig. 3a).

**Fig. 3.**
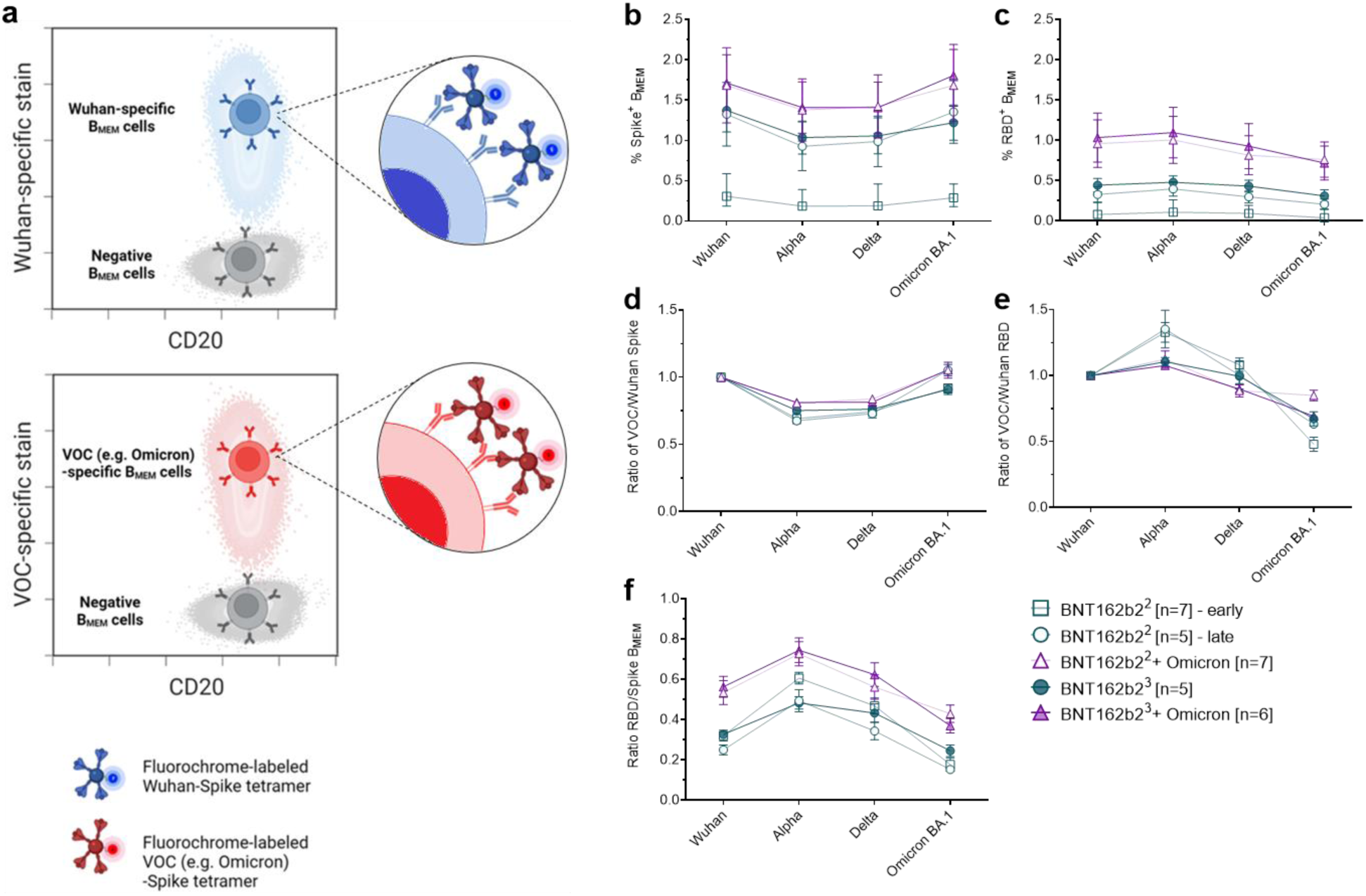
B_MEM_ cells of individuals double- and triple-vaccinated with BNT162b2 broadly recognize VOCs and are further boosted by Omicron breakthrough infection. Serum was drawn from double-vaccinated individuals (BNT162b2^2^) at 22 days after the second dose (green, open circles), from triple-vaccinated individuals (BNT162b2^3^) at 28 days after the third dose (green, closed circles), from double-vaccinated individuals with Omicron breakthrough infection (BNT162b2^2^ + Omi) at 46 days post-infection (purple, open triangles), and from triple-vaccinated individuals and Omicron breakthrough infection (BNT162b2^3^ + Omi) at 44 days post-infection (purple, closed triangles). Serum was tested in duplicate; 50% pseudovirus neutralization (pVN_50_) geometric mean titers (GMTs) (in a), 50% virus neutralization (VN_50_) GMTs (in b), and the geometric mean ratio of SARS-CoV-2 variant of concern (VOC) and Wuhan VN_50_ GMTs (in c) were plotted. For titer values below the limit of detection (LOD), LOD/2 values were plotted. Values above violin plots represent group GMTs. The non-parametric Friedman test with Dunn’s multiple comparisons correction was used to compare Wuhan neutralizing group GMTs with titers against the indicated variants and SARS-CoV-1. Multiplicity-adjusted p values are shown. (**a**) pVN_50_ GMTs against Wuhan, VOC and SARS-CoV-1 pseudovirus. (**b**) VN_50_ GMTs against live SARS-CoV-2 Wuhan, Beta, Delta and Omicron BA.1. (**c**) Group geometric mean ratios with 95% confidence intervals for all cohorts shown in b. PBMC samples from double-vaccinated individuals (BNT162b2^2^) at 22 days after the second dose (green, open squares) and 5 months after the second dose (green, open circles), from triple-vaccinated individuals (BNT162b2^3^) at 84 days after the third dose (green, closed circles), from double-vaccinated individuals with Omicron breakthrough infection (BNT162b2^2^ + Omi) at 46 days post-infection (purple, open triangles), and from triple-vaccinated individuals with Omicron breakthrough infection (BNT162b2^3^ + Omi) at 44 days post-infection (purple, closed triangles) were analyzed via flow cytometry for SARS-CoV-2-specific B_MEM_ cell (B_MEM_ – CD3-CD19+CD20+IgD-CD38^int/low^) frequencies via B cell bait staining. (**a**) Schematic of one-dimensional staining of B_MEM_ cells with fluorochrome-labeled SARS-CoV-2 S protein tetramer bait allowing discrimination of variant recognition. Frequencies of Wuhan or VOC full-length S protein-(**b**) and RBD-(**c**) specific B_MEM_ cells for the four groups of individuals were analyzed. Variant-specific B_MEM_ cell frequencies were normalized to Wuhan frequencies for S protein (**d**) and RBD-(**e**) binding. (**f**) Depicts the frequency ratios of RBD protein specific B_MEM_ cells over full-length S protein-specific B_MEM_ cells. Schematic in (a) was created with BioRender.com

As expected, the overall frequency of antigen-specific B_MEM_ cells varied across the different groups. Consistent with prior reports (*27*), the frequency of B_MEM_ cells in Omicron-naïve double-vaccinated individuals was low at an early time point after vaccination and increased over time: At 5 months as compared to 3 weeks after the second BNT162b2 dose, S protein-specific B_MEM_ cells almost quadrupled, RBD-specific ones tripled across all VOCs thereby reaching quantities similar to those observed in Omicron-naïve triple-vaccinated individuals (Fig. 3b, c, fig. S4a, b, c, Table S12).

Double or triple BNT162b2-vaccinated individuals with a SARS-CoV-2 Omicron breakthrough infection exhibited a strongly increased frequency of B_MEM_ cells, which was higher than those of Omicron-naïve triple-vaccinated individuals (Fig. 3b, d fig. S4d, e, k, l).

In all groups, including Omicron-naïve and Omicron infected individuals, B_MEM_ cells against Omicron BA.1 S protein were detectable at frequencies comparable to those against Wuhan and other tested VOCs (Fig 3b, d), whereas the frequency of B_MEM_ cells against Omicron BA.1 RBD was slightly lower compared to the other variants (Figure 3c, e, fig. S4f-j, m, n).

We then compared the ratios of RBD-to S protein-binding B_MEM_ cells within the different groups and found that they are biased towards S protein recognition for the Omicron BA.1 VOC, particularly in the Omicron-naïve groups (Figure 3f). In the Omicron-experienced groups this ratio is higher, indicating that an Omicron breakthrough infection improved Omicron BA.1 RBD recognition.

### Omicron breakthrough infection of BNT162b2 double- and triple-vaccinated individuals primarily boosts B_MEM_ cells against conserved epitopes shared broadly between S proteins of Wuhan and other VOCs rather than strictly Omicron S protein-specific epitopes

Our findings imply that Omicron infection in vaccinated individuals boosts not only neutralizing activity and B_MEM_ cells against Omicron BA.1, but broadly augments immunity against various VOCs. To investigate the specificity of antibody responses at a cellular level, we performed multi-parameter analyses of B_MEM_ cells stained with fluorescently labeled variant-specific S or RBD proteins.

By applying a combinatorial gating strategy, we sought to distinguish between B_MEM_ cell subsets that could identify only single variant-specific epitopes of Wuhan, Alpha, Delta or Omicron BA.1, versus those that could identify any given combination thereof (Fig 4a, fig. S3).

**Fig. 4.**
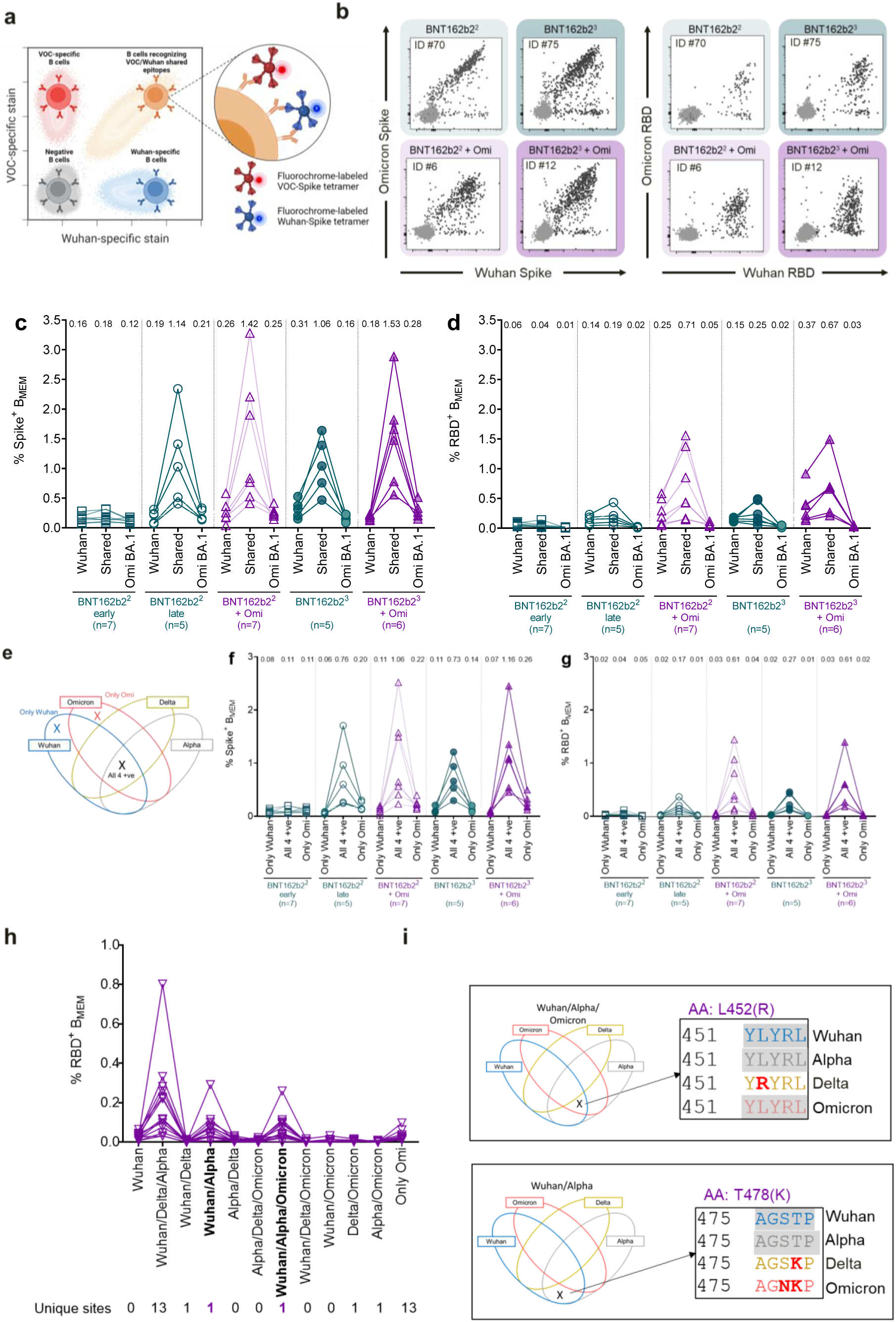
Omicron breakthrough infection of BNT162b2 double- and triple-vaccinated individuals primarily boosts B_MEM_ against conserved epitopes shared broadly between S proteins of Wuhan and other VOCs rather than strictly Omicron S-specific epitopes. PBMC samples from double-vaccinated individuals (BNT162b2^2^) at 22 days after the second dose (green, open squares) and 5 months after the second dose (green, open circles), from triple-vaccinated individuals (BNT162b2^3^) at 84 days after the third dose (green, closed circles), from double-vaccinated individuals with Omicron breakthrough infection (BNT162b2^2^ + Omi) at 46 days post-infection (purple, open triangles), and from triple-vaccinated individuals with Omicron breakthrough infection (BNT162b2^3^ + Omi) at 44 days post-infection (purple, closed triangle) were analyzed via flow cytometry for SARS-CoV-2-specific memory B cell (B_MEM_ – CD3-CD19+CD20+IgD-CD38^int/low^) frequencies via B cell bait staining (**a**). (**b**) Representative flow plots of Omicron and Wuhan S protein- and RBD-binding for each of the four groups of individuals investigated. Frequencies of B_MEM_ binding Omicron, Wuhan, or both (shared) full-length S protein (**c**) or RBD (**d**) for Omicron-experienced and naïve BNT162b2 double and triple vaccinees are shown. (**e**) Venn diagrams visualizing the combinatorial (Boolean) gating strategy to identify cross-reactive B_MEM_ recognizing all four variants simultaneously (All 4 +ve) and B_MEM_ recognizing only Omicron (only Omi) or only Wuhan (only Wuhan) S proteins. Frequencies for these three B_MEM_ sub-groups were compared for full-length S protein (**f**) and RBD (**g**) in the four different groups of individuals investigated. RBD variant recognition pattern by B_MEM_ was assessed through Boolean flow cytometric gating strategy and frequencies recognizing combinations of Wuhan and/or variant RBDs are displayed in (**h**), for all Omicron convalescent subjects (double and triple vaccinees pooled, n=13). (**i**) Conserved site within the RBD domain recognized by RBD-specific B_MEM_ after Omicron break-through infection. Mean values are indicated in c, d, f, and g. n = number of individuals per group. Schematic in (a) was created with BioRender.com

In a first analysis, we evaluated B_MEM_ cell recognition of Wuhan and Omicron BA.1 S and RBD proteins (Fig. 4b, c, d). The SARS-CoV-2 Omicron variant has 37 amino acid alterations in the S protein compared to the Wuhan parental strain, of which 15 alterations are in the RBD (*14*) (fig S5), an immunodominant target of neutralizing antibodies induced by COVID-19 vaccines or by SARS-CoV-2 infections (*15, 16*).

Staining with full length S proteins showed that the largest proportion of B_MEM_ cells from Omicron-naïve double-vaccinated individuals, and even more predominantly from triple-vaccinated individuals were directed against epitopes shared by both Wuhan and Omicron BA.1 SARS-CoV-2 variants. Consistent with the fact that vaccination with BNT162b2 can elicit immune responses against Wuhan epitopes that do not recognize the corresponding altered epitopes in the Omicron BA.1 S protein (Fig. 4b, c), we found in most individuals a smaller but clearly detectable proportion of B_MEM_ cells that recognized only Wuhan S protein or RBD. Consistent with the lack of exposure, no B_MEM_ cells binding exclusively to Omicron BA.1 S or RBD protein were detected in these Omicron-naïve individuals.

In Omicron convalescent individuals, frequencies of B_MEM_ cells recognizing S protein epitopes shared between Wuhan and Omicron BA.1 were significantly higher than in the Omicron-naïve ones (Fig 4b, c). In most of these subjects, we also found a small proportion of exclusively Wuhan S protein-specific B_MEM_ cells, as well as a slightly lower frequency of exclusively Omicron BA.1 variant S protein-specific ones.

A similar but slightly different pattern was observed by B cell staining with labeled RBD proteins (Fig 4b, d). Again, Omicron breakthrough infection of double-/triple-vaccinated individuals was found to primarily boost B_MEM_ cells reactive with conserved epitopes. A moderate boost of Wuhan-specific reactivities was observed; however, we could hardly detect only Omicron-RBD-specific B_MEM_ cells in the tested individuals (Fig. 4d).

Next, we employed the combinatorial gating approach to identify the subsets of S protein or RBD binding B_MEM_ cells that either bind exclusively to Wuhan or Omicron BA.1, or to common epitopes conserved broadly across all four variants, Wuhan, Alpha, Delta and Omicron BA.1 (Fig 4e). Across all four study groups, we found that the frequency of B_MEM_ cells recognizing S protein epitopes conserved across all tested variants accounted for the largest fraction of the pool of S protein-binding B_MEM_ cells (Fig. 4f, all 4+ve). The S protein of the Wuhan strain does not have an exclusive amino acid change that distinguishes it from the spike proteins of the Alpha, Delta, or Omicron BA.1 VOCs. Accordingly, we hardly detected B_MEM_ cells exclusively recognizing the Wuhan S protein in any individual (Fig. 4f). In several individuals with Omicron breakthrough infection, we detected a small proportion of B_MEM_ cells that bound exclusively to Omicron BA.1 S protein (Fig. 4f), whereas almost none of the individuals displayed a strictly Omicron BA.1 RBD-specific response (Fig. 4g).

Our findings indicate that SARS-CoV-2 Omicron breakthrough infection in vaccinated individuals primarily expands a broad B_MEM_ cell repertoire against conserved S protein and RBD epitopes rather than inducing large numbers of Omicron-specific B_MEM_ cells.

To further dissect the nuances of this response, we characterized the B_MEM_ subsets directed against the RBD. We used the combinatorial Boolean gating approach to discern B_MEM_ cells with distinct binding patterns in the spectrum of strictly variant-specific and common epitopes shared by several variants. Multiple sequence alignment revealed that the Omicron BA.1 RBD diverges from the RBD sequence regions conserved in Wuhan, Alpha and Delta by 13 single amino acid alterations (fig. S5). We found that all Omicron convalescent individuals had robust frequencies of B_MEM_ cells that recognized Wuhan, Alpha as well as the Delta VOC RBDs, but not Omicron BA.1 RBD, while B_MEM_ cells exclusively reactive with Omicron BA.1 RBD were almost absent in most of those individuals (Fig. 4h). We also did not detect B_MEM_ cells that exclusively recognized the Omicron BA.1 and Alpha RBDs, or the Omicron BA.1 and Delta RBDs.

Furthermore, in all individuals we identified two additional subsets of RBD-specific B_MEM_ cells. One subset was characterized by binding to Wuhan, Alpha as well as Omicron BA.1, but not the Delta, RBD. The other population exhibited binding to Wuhan and Alpha but not Omicron BA.1 or Delta RBD (Fig. 4h). Sequence alignment identified L452R as the only RBD mutation unique for Delta that is not shared by the other 3 variant RBDs (Fig. 4i top). Similarly, the only RBD site conserved in Wuhan and Alpha but altered in Delta and Omicron BA.1 was found to be T478K (Fig 4i bottom). Both L452R and T478K alterations are known to be associated with the evasion of vaccine induced neutralizing antibody responses (*29, 30*). Of note, no B_MEM_ cells were detected in all combinatorial subgroups in which multiple sequence alignment failed to identify unique epitopes in the RBD sequence that satisfied the Boolean selection criteria (e.g., Wuhan only or Wuhan and Omicron BA.1, but not Alpha, Delta). These observations indicate that the B_MEM_ cell response against RBD is driven by specificities induced through prior vaccination with BNT162b2 and not substantially redirected against new RBD epitopes mutated in the Omicron variant after infection.

## Conclusions

SARS-CoV-2 Omicron is a partial immune escape variant with an unprecedented number of amino acid alterations in the S protein at sites of neutralizing antibody binding, distinguishing it from previously reported variants (*14*). Recent neutralizing antibody mapping and molecular modeling studies strongly support the functional relevance of these alterations (*32, 33*), and their importance is confirmed by the fact that double-vaccinated individuals have no detectable neutralizing activity against SARS-CoV-2 Omicron (*22, 31*).

Our findings show that Omicron breakthrough infection of vaccinated individuals boosts not only neutralizing activity and B_MEM_ cells against Omicron but broadly augments immunity against various VOCs.

Our study also provides insights into how this broad immunity is achieved. Our data suggest that initial exposure to the Wuhan strain S protein may have shaped the formation of B_MEM_ cells and imprinted against the formation of novel B_MEM_ cell responses against the more distinctive epitopes of the Omicron variant. Similar observations have been reported from vaccinated individuals who experienced breakthrough infections with the delta variant (3). In conclusion, Omicron breakthrough infection primarily expands a broad B_MEM_ cell repertoire against conserved S protein and RBD epitopes, rather than inducing considerable numbers of strictly Omicron-specific B_MEM_ cells.

Thus, Omicron breakthrough infection in double-vaccinated individuals leads to expansion of the pre-existing B_MEM_ cell pool, similar to a third dose of booster vaccination. However, there are clear differences in the immune response pattern induced by a homologous vaccine booster as compared to an Omicron breakthrough infection. Despite the obvious focus of the B cell memory response on conserved epitopes, Omicron breakthrough infection leads to a more substantial increase in antibody neutralization titers against Omicron, as well as pronounced cross-neutralization of both the ancestral and the novel SARS CoV-2 variants. These effects are particularly striking in double-vaccinated individuals.

Three findings may point to potentially complementary and synergistic mechanisms responsible for these results:

First, an overall increase of S protein-specific B_MEM_ cells. Omicron-convalescent double-vaccinated individuals have a higher frequency of B_MEM_ cells and higher neutralizing antibody titers against all VOCs as compared to triple-vaccinated individuals. That breakthrough infection elicits a stronger neutralizing antibody response than the third vaccine dose in double-vaccinated individuals is not apparent from previous studies describing breakthrough infections with other variants (*4, 34*) and may be explained by poor neutralization of the Omicron variant in the initial phase of infection, potentially causing a greater or prolonged antigen exposure of the immune system to the altered S protein.

Second, a stronger bias on RBD-specific B_MEM_ cell responses. Omicron breakthrough infection promotes proportionally more pronounced boosting of RBD-specific B_MEM_ cells than of B_MEM_ cells that recognize S protein-specific epitopes outside the RBD. Therefore, Omicron-infected individuals have a significantly higher ratio of RBD/S protein-specific B_MEM_ cells compared to vaccinated Omicron-naïve individuals. The RBD is the key domain of the S protein that binds to the SARS-CoV-2 receptor ACE2 and harbors multiple neutralizing antibody binding sites including some regions that are not affected by Omicron alterations, such as those at position L452. An increased focus of the immune response on this domain could help restore immune protection by stimulating B_MEM_ cells that produce neutralizing antibodies against RBD epitopes unaltered in Omicron.

Third, the induction of broadly neutralizing antibodies. We found that the majority of sera from Omicron-convalescent but not from Omicron-naïve vaccinated individuals robustly neutralized SARS-CoV-1. This may suggest that Omicron infection in vaccinated individuals stimulates B_MEM_ cells that form neutralizing antibodies against spike protein epitopes conserved in the SARS-CoV-1 and SARS-CoV-2 families. Broadly neutralizing antibodies have recently been described in SARS-CoV-1 infected individuals vaccinated with BNT162b2 (*24*). Such pan-Sarbecovirus immune responses are thought to be triggered by neutralizing antibodies to highly conserved S protein domains (*35, 36*). The greater antigenic distance of the Omicron spike protein from the other SARS-Cov-2 strains may promote targeting of conserved subdominant neutralizing epitopes as recently described to be located in the C-terminal portion of the spike protein (*37, 38*). Future studies mapping monoclonal antibodies derived from Omicron-specific B_MEM_ cells will provide further insight into the relevance of these findings.

In aggregate, our results suggest that despite possible imprinting of the immune response by previous vaccination, the preformed B-cell memory pool can be refocused and quantitatively remodeled by exposure to heterologous S proteins to allow neutralization of variants that evade a previously established neutralizing antibody response.

Limitations of this study are that the data were generated from blood samples obtained a few weeks after Omicron infection. Later time points might reveal formation of memory responses against novel epitopes that are not yet visible. Also, the analysis presented here has evaluated sample sets from multiple studies with a limited sample size, different sampling time points and baseline or demographic characteristics. The analysis was limited to B_MEM_ cells while long-lived bone marrow-derived plasma cells (BMPCs) which are known to be BNT162b2 vaccination induced (*28*), could not be investigated as they cannot be cryopreserved.

In conclusion, while the data are based on samples from individuals exposed to the Omicron S protein as a result of infection, our observations suggest that a vaccine adapted to the Omicron S protein could similarly reshape the B-cell memory repertoire and therefore may be more beneficial than an extended series of boosters with the existing Wuhan-Hu-1 spike based vaccines.

## Acknowledgments

We thank the BioNTech German clinical Phase 1/2 trial (NCT04380701, EudraCT: 2020-001038-36), the German Phase 2 rollover booster trial (NCT04949490, EudraCT: 2021-002387-50) the global clinical Phase 2 trial (NCT04380701) participants, and the Omicron convalescent Research Study participants from whom the post-immunization human sera and PBMCs were obtained. We thank the many colleagues at BioNTech and Pfizer who developed and produced the BNT162b2 vaccine candidate. We thank Sabrina Jägle and Nina Beckmann for logistical support. We thank the VisMederi team for performing excellent work on live virus-neutralising antibody assays. We thank Svetlana Shpyro, Sayeed Nadim, Christina Heiser, Ayca Telorman, Claudia Müller, Amy Wanamaker, Nicki Williams and Jennifer VanCamp for sample demographics support.

## Author contributions

U.S., Ö.T., J.Q., and A.M. conceived and conceptualized the work. J.Q., A.M, B.G.L. and I.V. planned and supervised experiments. K.K., O.O., S.H., U.G. and S.C. coordinated and conducted sample collection. K.G. coordinated sample shipments and clinical data transfer. J.Q., A.M., B.G.L., S.L., A.W., P.A.., and M.B. performed experiments. J.Q., A.M., N.S., B.G.L., S.L., K.K., and O.O. analyzed data. U.S., Ö.T., J.Q., A.M., N.S., and A.F. interpreted data and wrote the manuscript. All authors supported the review of the manuscript.

## Data and materials availability

Participant baseline characteristics are provided in Table S1 to Table S3 and Table S10. The neutralization titers are provided in Tables S4 to S9 and S11. The frequencies of SARS-CoV-2 variant S protein/RBD-specific B_MEM_ cells are provided in Table S12. Materials are available from the authors under a material transfer agreement with BioNTech.

## Supplementary Materials

Materials and Methods

Figs. S1-S6

Tables S1-S12

