## Supplement for "Omicron breakthrough infection drives cross-variant neutralization and memory B cell formation"

**This PDF file includes:**

Materials and Methods

Fig. S1 to S6

Tables S1 to S12

References (for supplementary section)

### Materials and Methods

Recruitment of participants and sample collection

Individuals from the SARS-CoV-2 Omicron-naïve BNT162b2 double-vaccinated ($BNT162b2^2$) and triple-vaccinated ($BNT162b2^3$) cohorts provided informed consent as part of their participation in a clinical trial (the Phase 1/2 trial BNT162-01 [NCT04380701] (*25*), the Phase 2 rollover trial BNT162-14 [NCT04949490], or as part of the BNT162-17 [NCT05004181] trial). Participants from the SARS-CoV-2 Omicron convalescent double- and triple vaccinated cohorts ($BNT162b2^2$ + Omi and $BNT162b2^3$ + Omi cohorts, respectively) and individuals vaccinated with other approved COVID-19 vaccines or mixed regimens with subsequent Omicron breakthrough infection were recruited from University Hospital, Goethe University Frankfurt as part of a research program that recruited patients that had experienced Omicron breakthrough infection following vaccination for COVID-19, to provide blood samples and clinical data for research. Infection with the Omicron strain was confirmed with variant-specific PCR or sequencing, and participants were free of symptoms at the time of blood collection. The study protocol for this research program was approved by the Ethics Board of the University Hospital, Goethe University Frankfurt (No. 2021-560).

Demographic and clinical information for all participants as well as sampling timepoints are provided in Tables S1-S3 and S10, and Fig. 1.

Serum was isolated by centrifugation 2000 x g for 10 minutes and cryopreserved until use. Li-Heparin blood samples were isolated by density gradient centrifugation using Ficoll-Paque PLUS (Cytiva) and were subsequently cryopreserved until use.

VSV-SARS-CoV-2 S variant pseudovirus generation

A recombinant replication-deficient vesicular stomatitis virus (VSV) vector that encodes green fluorescent protein (GFP) and luciferase instead of the VSV-glycoprotein (VSV-G) was pseudotyped with SARS-CoV-1 spike (S) (UniProt Ref: P59594) and with SARS-CoV-2 S derived from either the Wuhan reference strain (NCBI Ref: 43740568), the Alpha variant (mutations: Δ69/70, Δ144, N501Y, A570D, D614G, P681H, T716I, S982A, D1118H), the Beta variant (mutations: L18F, D80A, D215G, Δ242–244, R246I, K417N, E484K, N501Y, D614G, A701V), the Delta variant (mutations: T19R, G142D, E156G, Δ157/158, K417N, L452R, T478K, D614G, P681R, D950N) the Omicron BA.1 variant (mutations: A67V, Δ69/70, T95I, G142D, Δ143-145, Δ211, L212I, ins214EPE, G339D, S371L, S373P, S375F, K417N, N440K, G446S, S477N, T478K, E484A, Q493R, G496S, Q498R, N501Y, Y505H, T547K, D614G, H655Y, N679K, P681H, N764K, D796Y, N856K, Q954H, N969K, L981F) or the Omicron BA.2 variant (mutations: T19I, Δ24-26, A27S, G142D, V213G, G339D, S371F, S373P, S375F, T376A, D405N, R408S, K417N, N440K, S477N, T478K, E484A, Q493R, Q498R, N501Y, Y505H, D614G, H655Y, N679K, P681H, N764K, D796Y, Q954H, N969K) according to published pseudotyping protocols (*39*).

A diagram of SARS-CoV-2 S protein mutations is shown in fig. S6a. In brief, HEK293T/17 monolayers (ATCC® CRL-11268™) cultured in Dulbecco's modified Eagle's medium (DMEM) with GlutaMAX™ (Gibco) supplemented with 10% heat-inactivated fetal bovine serum (FBS [Sigma-Aldrich]) (referred to as medium) were transfected with Sanger sequencing-verified SARS-CoV-1 or variant-specific SARS-CoV-2 S expression plasmid with Lipofectamine LTX (Life Technologies) following the manufacturer's instructions. At 24 hours VSV-G complemented VSVΔG vector. After incubation for 2 hours at 37 °C with 7.5% $CO_2$, cells were washed twice with phosphate buffered saline (PBS) before medium supplemented with anti-VSV-G antibody (clone 8G5F11, Kerafast Inc.) was added to neutralize residual VSV-G-complemented input virus. VSV-SARS-CoV-2-S pseudotype-containing medium was harvested 20 hours after inoculation, passed through a 0.2 µm filter (Nalgene) and stored at -80 °C. The pseudovirus batches were titrated on Vero 76 cells (ATCC® CRL-1587™) cultured in medium. The relative luciferase units induced by a defined volume of a Wuhan spike pseudovirus reference batch previously described in Muik et al., 2021 (*23*), that corresponds to an infectious titer of 200 transducing units (TU) per mL, was used as a comparator. Input volumes for the SARS-CoV-2 variant pseudovirus batches were calculated to normalize the infectious titer based on the relative luciferase units relative to the reference.

Pseudovirus neutralization assay

Vero 76 cells were seeded in 96-well white, flat-bottom plates (Thermo Scientific) at 40,000 cells/well in medium 4 hours prior to the assay and cultured at 37 °C with 7.5% $CO_2$. Each serum was serially diluted 2-fold in medium with the first dilution being 1:5 (Omicron-naïve double- and triple BNT162b2 vaccinated; dilution range of 1:5 to 1:5,120) or 1:30 (double- and triple BNT162b2 vaccinated after subsequent Omicron breakthrough infection; dilution range of 1:30 to 1:30,720). VSV-SARS-CoV-2-S/VSV-SARS-CoV-1-S particles were diluted in medium to obtain 200 TU in the assay. Serum dilutions were mixed 1:1 with pseudovirus (n=2 technical replicates per serum per pseudovirus) for 30 minutes at room temperature before being added to Vero 76 cell monolayers and incubated at 37 °C with 7.5% $CO_2$ for 24 hours. Supernatants were removed and the cells were lysed with luciferase reagent (Promega). Luminescence was recorded on a CLARIOstar® Plus microplate reader (BMG Labtech), and neutralization titers were calculated as the reciprocal of the highest serum dilution that still resulted in 50% reduction in luminescence. Results were expressed as geometric mean titers (GMT) of duplicates. If no neutralization was observed, an arbitrary titer value of half of the limit of detection [LOD] was reported. Tables of the neutralization titers are provided (Table S4-6 and Table S11).

Live SARS-CoV-2 neutralization assay

SARS-CoV-2 virus neutralization titers were determined by a microneutralization assay based on cytopathic effect (CPE) at VisMederi S.r.l., Siena, Italy. In brief, heat-inactivated serum samples from participants were serially diluted 1:2 (starting at 1:10) and incubated for 1 hour at 37 °C with 100 $TCID_{50}$ of live Wuhan-like SARS-CoV-2 virus strain 2019-nCOV/ITALY-INMI1 (GenBank: MT066156), Beta virus strain Human nCoV19 isolate/England ex-SA/HCM002/2021 (mutations: D80A, D215G, Δ242–244, K417N, E484K, N501Y, D614G, A701V), sequence-verified Delta strain isolated from a nasopharyngeal swab (mutations: T19R, G142D, E156G, Δ157/158, L452R, T478K, D614G, P681R, R682Q, D950N) or Omicron BA.1 strain hCoV-19/Belgium/rega-20174/2021 (mutations: A67V, Δ69/70, T95I, G142D, Δ143-145, Δ211, L212I, ins214EPE, G339D, S371L, S373P, S375F, K417N, N440K, G446S, S477N, T478K, E484A, Q493R, G496S, Q498R, N501Y, Y505H, T547K, D614G, H655Y, N679K, P681H, N764K, D796Y, N856K, Q954H, N969K, L981F) to allow any antigen-specific antibodies to bind to the virus. A diagram of S protein mutations is shown in fig. S6b. The 2019-nCOV/ITALY-INMI1 strain S protein is identical in sequence to the wild-type SARS-CoV-2 S (Wuhan-Hu-1 isolate). Vero E6 (ATCC® CRL-1586™) cell monolayers were inoculated with the serum/virus mix in 96-well plates and incubated for 3 days (2019-nCOV/ITALY-INMI1 strain) or 4 days (Beta,

Delta and Omicron BA.1 variant strain) to allow infection by non-neutralized virus. The plates were observed under an inverted light microscope and the wells were scored as positive for SARS-CoV-2 infection (i.e., showing CPE) or negative for SARS-CoV-2 infection (i.e., cells were alive without CPE). The neutralization titer was determined as the reciprocal of the highest serum dilution that protected more than 50% of cells from CPE and reported as GMT of duplicates. If no neutralization was observed, an arbitrary titer value of 5 (half of the LOD) was reported. Tables of the neutralization titers are provided (Table S7 to S9).

Detection and characterization of SARS-CoV-2-specific B cells with flow cytometry

Spike (S) protein/Receptor-binding domain (RBD)-specific B cells were detected using recombinant, biotinylated SARS-CoV-2 Spike (Acro Biosystems: Wuhan – SPN-C82E9, Alpha – SPN-C82E5, Delta – SPN-C82Ec, Omicron – SPN-C82Ee) and RBD (Acro Biosystems: Wuhan – SPD-B28E9, Alpha – SPD-C82E6, Delta – SPD-C82Ed, Omicron – SPD-C82E4) proteins. Recombinant Spike and RBD proteins were tetramerized with fluorescently labeled Streptavidin (BioLegend, BD Biosciences) in a 4:1 molar ratio for 1 h at 4 °C in the dark. Afterwards samples were spun down for 10 min at 4°C to remove eventual precipitates.

For flow cytometric analysis, PBMCs were thawed and $5x10^6$ cells per sample were seeded into 96 U-bottom plates. Cells were blocked for Fc-receptor-binding (Human BD Fc Block™, BD Biosciences) and statured with free Biotin (D-Biotin, Invitrogen, 1 µM) in flow buffer (DPBS (Gibco) supplemented with 2% FBS (Sigma), 2 mM EDTA (Sigma-Aldrich)) for 20 min at 4 °C. Cells were washed and labeled with BCR bait tetramers supplemented with free Biotin in flow buffer (D-Biotin, Invitrogen, 2 µg/ml) for 1 h at 4 °C in the dark (2 µg/ml for Spike and 0,25 µg/ml for RBD proteins). Cells were washed with flow buffer and stained for viability (Fixable

Viability Dye eFluor™ 780, eBioscience) and surface markers (CD3 – clone: UCHT1(BD Biosciences), CD4 – clone: SK3 (BD Biosciences), CD185 (CXCR5) – clone: RF8B2 (BioLegend), CD279 (PD-1) – clone: EH12.1(BD Biosciences), CD278 (ICOS) – clone: C398.4A (BioLegend), CD19- clone: SJ25C1(BD Biosciences), CD20 – clone: 2H7(BD Biosciences), CD21 – clone: B-ly4(BD Biosciences), CD27 – clone: L128(BD Biosciences), CD38 – clone: HIT2(BD Biosciences), CD11c – clone: S-HCL-3(BD Biosciences), CD138 – clone: MI15(BD Biosciences), IgG - clone: G18-145(BD Biosciences), IgM – clone: G20-127(BD Biosciences), IgD – clone: IA6-2(BD Biosciences), CD14 – clone: MφP9 (BD Biosciences, dump channel), CD16 – clone: 3G8 (BD Biosciences, dump channel)) in flow buffer supplemented with Brilliant Stain Buffer Plus (BD Biosciences, according to the manufacturer's instructions) for 20 min at 4 °C. Samples were washed and fixed with BD™ Stabilizing Fixative (BD Biosciences, according to the manufacturer's instructions) prior to data acquisition on a BD Symphony A3 flow cytometer. FCS 3.0 files were exported from BD Diva Software and analyzed using FlowJo software (Version 10.7.1.).

Statistical analysis

The statistical method of aggregation used for the analysis of antibody titers is the geometric mean and for the ratio of SARS-CoV-2 VOC titer and Wuhan titer the geometric mean and the corresponding 95% confidence interval. The use of the geometric mean accounts for the non-normal distribution of antibody titers, which span several orders of magnitude. The Friedman test with Dunn's correction for multiple comparisons was used to conduct pairwise signed-rank tests of group geometric mean neutralizing antibody titers with a common control group. Flow cytometric frequencies were analyzed with and tables were exported from FlowJo software

(Version 10.7.1.). Statistical analysis of cumulative memory B cell frequencies was the mean and standard errors of the mean (SEM). All statistical analyses were performed using GraphPad Prism software version 9.

### Fig. S1

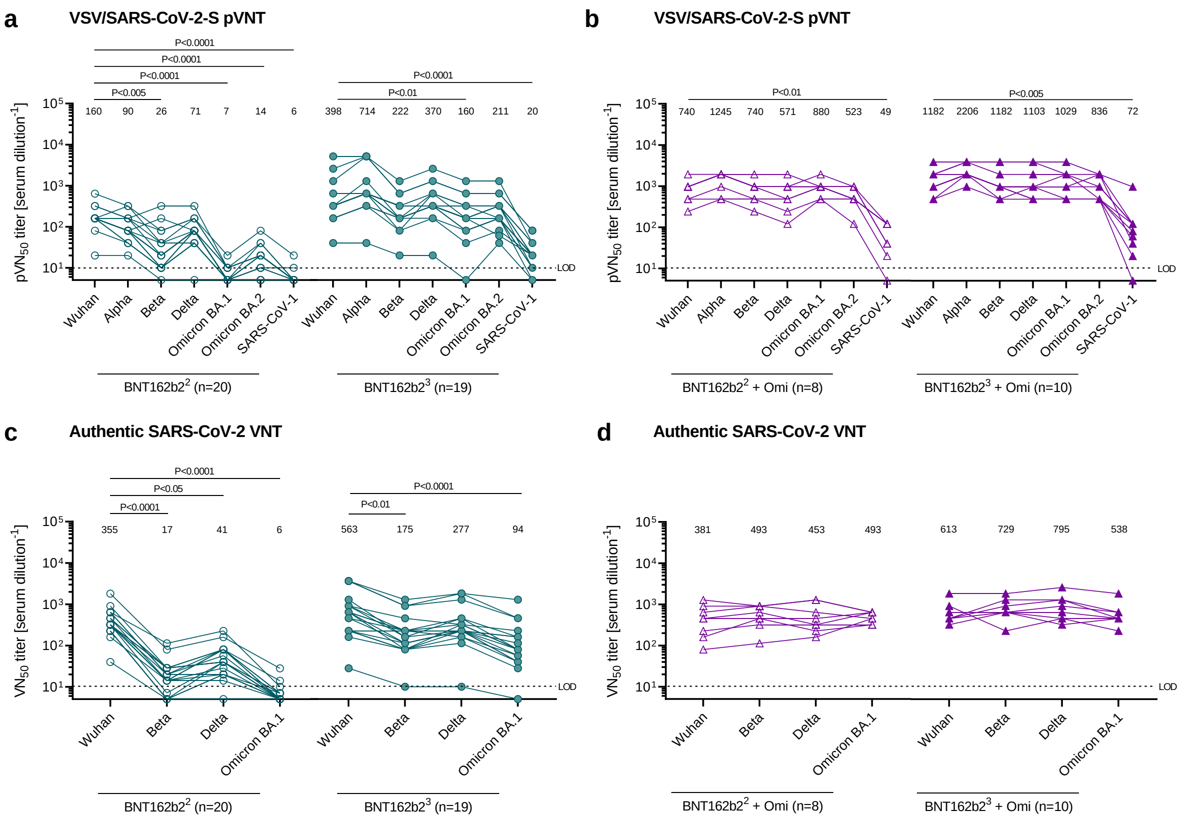


**Fig. S1. Individually plotted 50% pseudovirus and live SARS-CoV-2 virus neutralization titers (pVN50 and VN50) against SARS-CoV-2 Wuhan, variants of concern (VOCs) and SARS-CoV-1.**

Individual line plots from the dataset shown in Figure 2. pVN50 GMTs against Wuhan, VOC and SARS-CoV-1 pseudovirus are shown for (**a**) Omicron-naïve individuals double- (BNT162b2$^2$: green, open circles) and triple-vaccinated with BNT162b2 (BNT162b2$^3$: green, closed circles), and (**b**) Omicron breakthrough infected individuals double- (BNT162b22 + Omi: purple, open triangles) and triple-vaccinated with BNT162b2 (BNT162b2$^3$ + Omi: purple, closed triangles) prior to infection. VN50 GMTs against Wuhan and VOC are shown for (**c**) Omicron-naïve individuals double- (BNT162b2$^2$: green, open circles) and triple-vaccinated with BNT162b2 (BNT162b2$^3$: green, closed circles), and (**d**) Omicron breakthrough infected individuals double- (BNT162b2$^2$ + Omi: purple, open triangles) and triple-vaccinated with BNT162b2 (BNT162b2$^3$ + Omi: purple, closed triangles) prior to infection. Serum was tested in duplicate. For titer values below the LOD, LOD/2 values were plotted. Values above line plots represent group GMTs. The non-parametric Friedman test with Dunn's multiple comparisons correction was used to compare Wuhan neutralizing group GMTs with titers against the indicated variants and SARS-CoV-1. Multiplicity-adjusted p values are shown.

**Fig. S2**

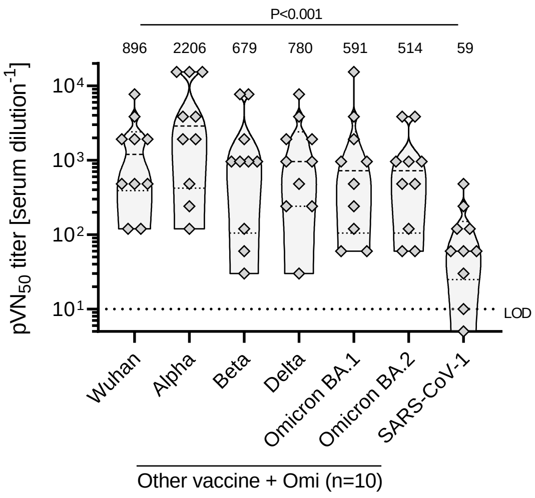


**Fig. S2. Omicron breakthrough infection of individuals vaccinated with other approved COVID-19 vaccines or mixed regimens results in immune sera that broadly neutralize Omicron BA.1, BA.2 and other VOCs plus SARS-CoV-1**

Serum was drawn from 10 individuals vaccinated with other approved COVID-19 vaccines or mixed regimens at a median 43 days after infection (grey diamonds). Serum was tested in duplicate; individual 50% pseudovirus neutralization (pVN50) geometric mean titers (GMTs) against SARS-CoV-2 Wuhan, Alpha, Beta, Delta and Omicron BA.1 and BA.2 variants, plus SARS-CoV-1 were plotted. For titer values below the limit of detection (LOD), LOD/2 values were plotted. Values above violin plots represent group GMTs. The non-parametric Friedman test with Dunn's multiple comparisons correction was used to compare Wuhan neutralizing group GMTs with titers against the indicated variants and SARS-CoV-1. Multiplicity-adjusted p values are shown.

### Fig. S3

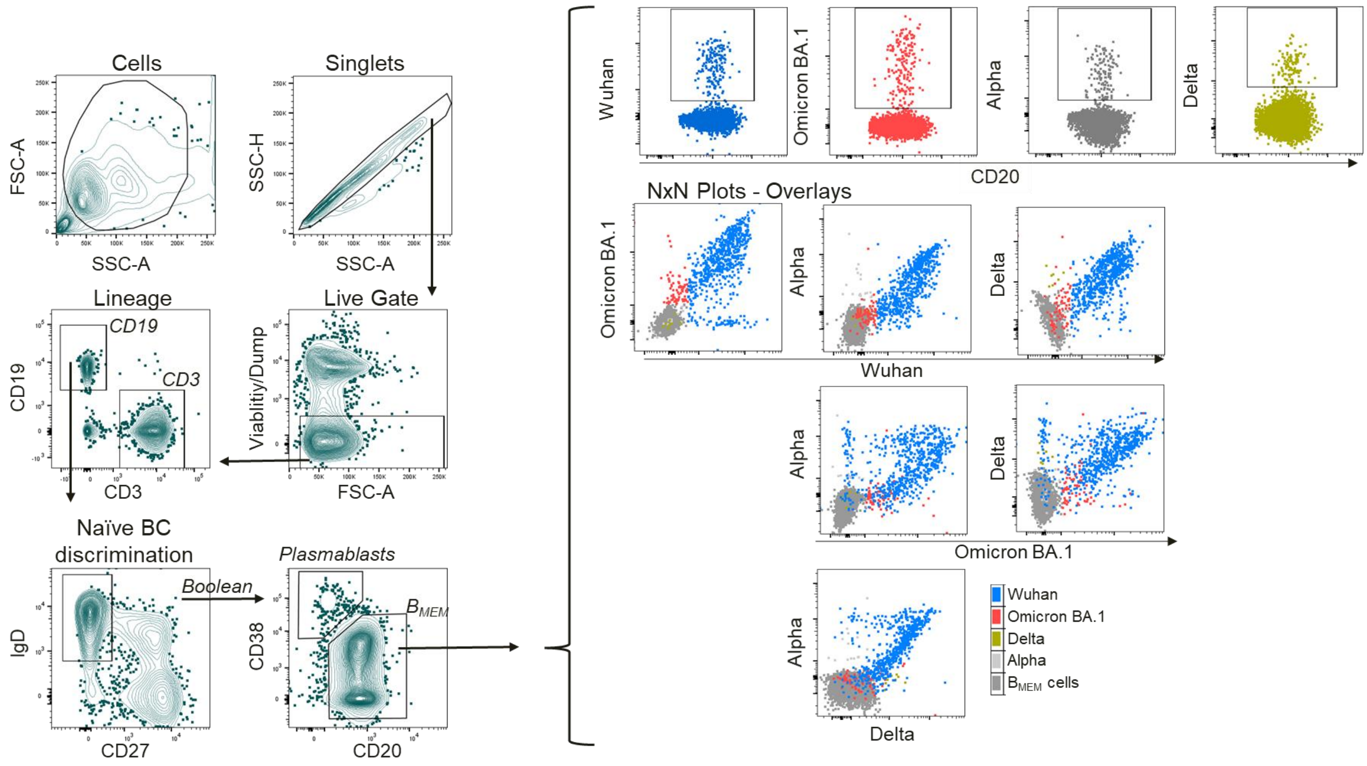


**Fig. S3. Flow cytometric gating strategy for B cell phenotypic analysis.**

Representative flow plots of a triple BNT162b2 vaccinated and Omicron convalescent subject analyzed for binding of full-length SARS-CoV-2 Wuhan Spike protein and its variants (also referred to as B cell baits). Debris and doublets were discriminated via FSC/SSC. Then dead cells and monocytes (CD14, CD16 – Viability/Dump channel) were excluded. CD19 positive B cells were analyzed for IgD and CD27 expression, thereby naïve B cells were discriminated as $IgD^{+}$ cells with the Boolean 'make non-gate' function. Within non-naïve B cells Plasmablasts ($CD38^{high}$ $CD20^{low}$) and memory B cells ($B_{MEM}$s $CD38^{int/low}CD20^{high}$) were distinguished. $B_{MEM}$ cells were analyzed for B cell bait binding. SARS-CoV-2 Spike reactivities were assessed by gating on each Spike/RBD variant tested by plotting against the CD20 signal. Bait gates were overlayed onto total $B_{MEM}$ cells and displayed as NxN-Plots for the four bait channels.

**Fig. S4**

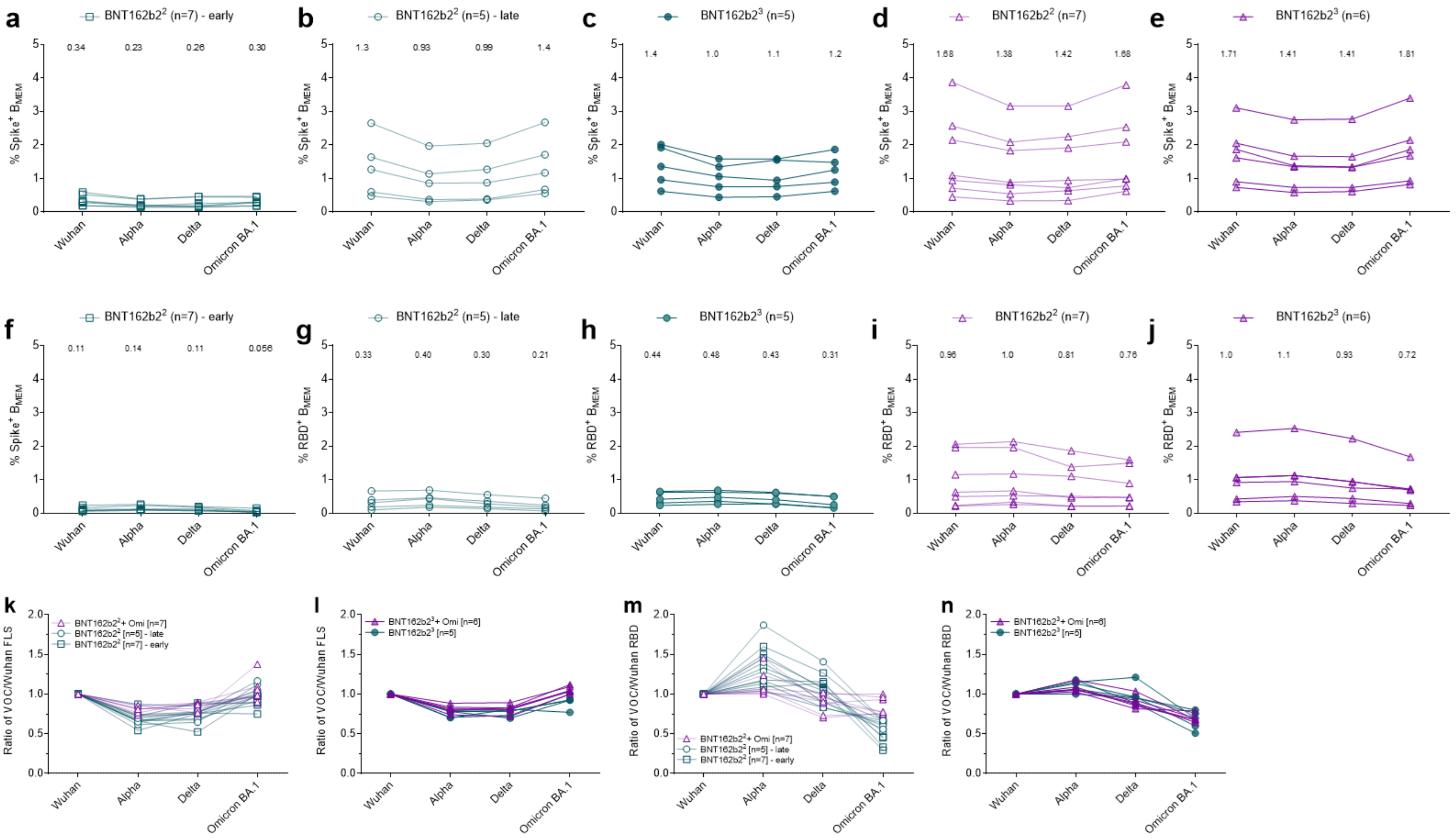

a
BNT162b2² (n=7) - early
0.34
0.23
0.26
0.30
% Spike⁺ B_MEM
Wuhan
Alpha
Delta
Omicron BA.1
b
BNT162b2² (n=5) - late
1.3
0.93
0.99
1.4
c
BNT162b2³ (n=5)
1.4
1.0
1.1
1.2
d
BNT162b2² (n=7)
1.68
1.38
1.42
1.68
e
BNT162b2³ (n=6)
1.71
1.41
1.41
1.81
f
BNT162b2² (n=7) - early
0.11
0.14
0.11
0.056
g
BNT162b2² (n=5) - late
0.33
0.40
0.30
0.21
% RBD⁺ B_MEM
h
BNT162b2³ (n=5)
0.44
0.48
0.43
0.31
i
BNT162b2² (n=7)
0.96
1.0
0.81
0.76
j
BNT162b2³ (n=6)
1.0
1.1
0.93
0.72
k
BNT162b2²+ Omi [n=7]
BNT162b2² [n=5] - late
BNT162b2² [n=7] - early
Ratio of VOC/Wuhan FLS
l
BNT162b2³+ Omi [n=6]
BNT162b2³ [n=5]
m
Ratio of VOC/Wuhan RBD
n

**Fig. S4. Omicron breakthrough infection of BNT162b2 double- and of triple-vaccinated individuals primarily boosts $B_{MEM}$ against conserved epitopes shared broadly between S proteins of Wuhan and other VOCs rather than strictly Omicron spike-specific epitopes.**

PBMC samples from double (BNT162b2$^2$) and triple (BNT162b2$^3$) BNT162b2 vaccinated individuals who did (green) or did not (purple) experience an Omicron breakthrough infection (+Omi) were analyzed via flow cytometry for antigen-specific memory B cells ($B_{MEM}$ – $CD3^-CD19^+CD20^+IgD^-CD38^{int/low}$) frequencies via B cell bait staining. Frequencies of full-length of Wuhan, Alpha, Delta, and Omicron S specific $B_{MEM}$ (**a-e**), and Wuhan and variants RBD-specific $B_{MEM}$ cells (**f-j**) for the five different groups of individuals are shown. Each line represents one individual donor. Variant-specific $B_{MEM}$ frequencies were normalized to Wuhan frequencies for full-length S (**k, l**) and RBD (**m, n**) binding for Omicron-naïve and -experienced individuals that received to doses of BNT162b2 (**k, m**) and triple dosed individuals (**l, n**). Each line represents one individual donor. Mean values are indicated. n = number of individuals per group.

### Fig. S5

```
RBD
Wuhan
VOC ALPHA
VOC DELTA
VOC OMICRON BA.1

301  CTLKSFTVEKGIYQTSNFRVQPTESIVRFPNITNLCPFGEVFNATRFASV
301  CTLKSFTVEKGIYQTSNFRVQPTESIVRFPNITNLCPFGEVFNATRFASV
301  CTLKSFTVEKGIYQTSNFRVQPTESIVRFPNITNLCPFGEVFNATRFASV
301  CTLKSFTVEKGIYQTSNFRVQPTESIVRFPNITNLCPFDEVFNATRFASV

351  YAWNRKRISNCVADYSVLYNSASFSTFKCYGVSPTKLNDLCFTNVYADSF
351  YAWNRKRISNCVADYSVLYNSASFSTFKCYGVSPTKLNDLCFTNVYADSF
351  YAWNRKRISNCVADYSVLYNSASFSTFKCYGVSPTKLNDLCFTNVYADSF
351  YAWNRKRISNCVADYSVLYNLAPFFTFKCYGVSPTKLNDLCFTNVYADSF

401  VIRGDEVRQIAPGQTGKIADYNYKLPDDFTGCVIAWNSNNLDSKVGGNYN
401  VIRGDEVRQIAPGQTGKIADYNYKLPDDFTGCVIAWNSNNLDSKVGGNYN
401  VIRGDEVRQIAPGQTGKIADYNYKLPDDFTGCVIAWNSNNLDSKVGGNYN
401  VIRGDEVRQIAPGQTGNIADYNYKLPDDFTGCVIAWNSNKLDSKVSGNYN

451  YLYRLFRKSNLKPFERDISTEIYQAGSTPCNGVEGFNCYFPLQSYGFQPT
451  YLYRLFRKSNLKPFERDISTEIYQAGSTPCNGVEGFNCYFPLQSYGFQPT
451  YRYRLFRKSNLKPFERDISTEIYQAGSKPCNGVEGFNCYFPLQSYGFQPT
451  YLYRLFRKSNLKPFERDISTEIYQAGNKPCNGVAGFNCYFPLRSYSFRPT

501  NGVGYQPYRVVVLSFELLHAPATVCGPKKSTNLVKNKCVNFNFNGLTGTG
501  YGVGYQPYRVVVLSFELLHAPATVCGPKKSTNLVKNKCVNFNFNGLTGTG
501  NGVGYQPYRVVVLSFELLHAPATVCGPKKSTNLVKNKCVNFNFNGLTGTG
501  YGVGHQPYRVVVLSFELLHAPATVCGPKKSTNLVKNKCVNFNFNGLKGTG
```

**Fig. S5. Sequences of Wuhan and variants RBDs.**

Variant-specific point-mutations are indicated in bold red font.

**Fig. S6**

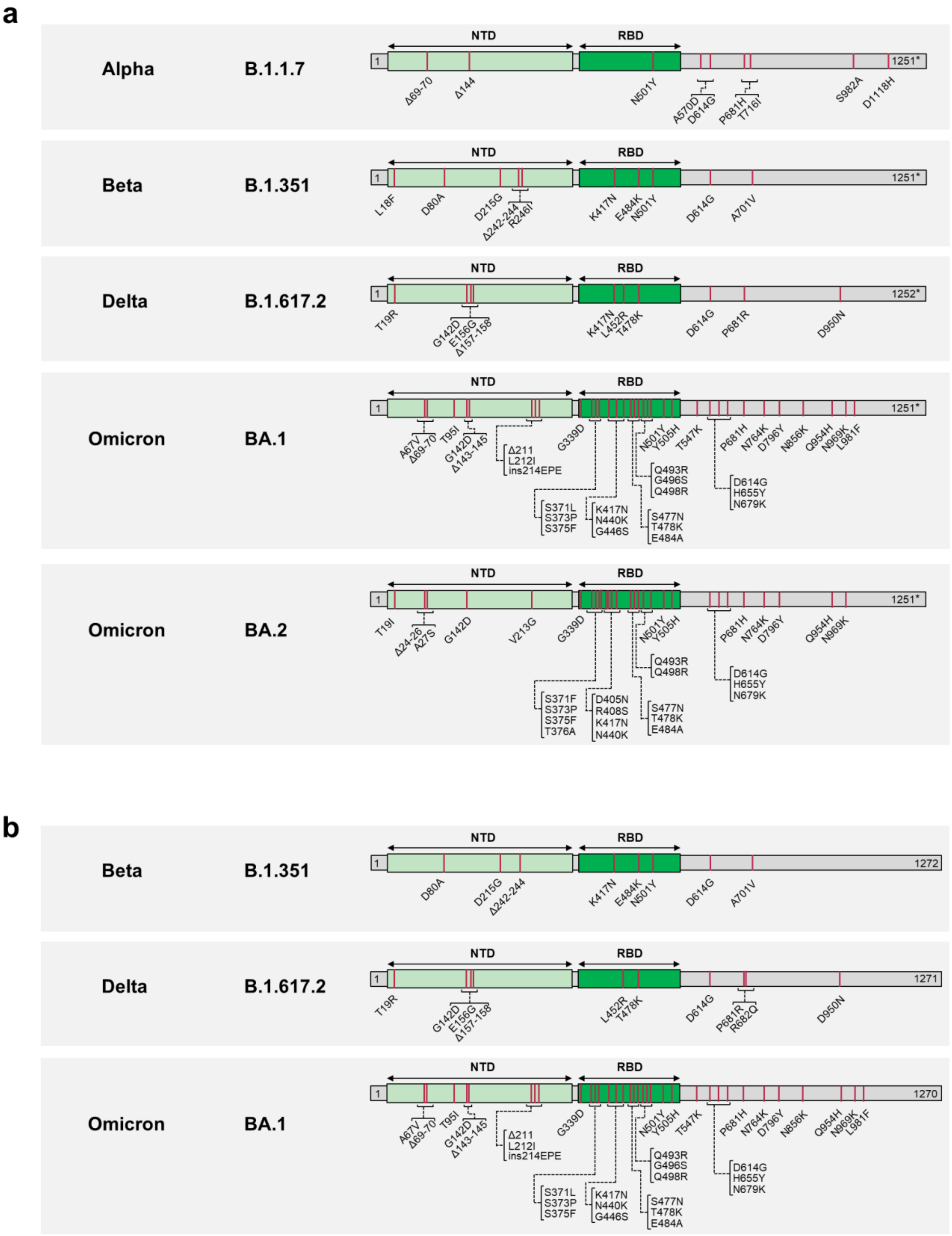
a
Alpha
B.1.1.7
NTD
RBD
1
1251*
Δ69-70
Δ144
N501Y
A570D
D614G
P681H
T716I
S982A
D1118H
Beta
B.1.351
L18F
D80A
D215G
Δ242-244
R246I
K417N
E484K
N501Y
D614G
A701V
Delta
B.1.617.2
1252*
T19R
G142D
E156G
Δ157-158
K417N
L452R
T478K
D614G
P681R
D950N
Omicron
BA.1
A67V
Δ69-70
T95I
G142D
Δ143-145
Δ211
L212I
ins214EPE
G339D
S371L
S373P
S375F
K417N
N440K
G446S
S477N
T478K
E484A
Q493R
G496S
Q498R
N501Y
Y505H
T547K
D614G
H655Y
N679K
P681H
N764K
D796Y
N856K
Q954H
N969K
L981F
Omicron
BA.2
T19I
Δ24-26
A27S
G142D
V213G
G339D
S371F
S373P
S375F
T376A
D405N
R408S
K417N
N440K
S477N
T478K
E484A
Q493R
Q498R
N501Y
Y505H
D614G
H655Y
N679K
P681H
N764K
D796Y
Q954H
N969K
b
Beta
B.1.351
1272
D80A
D215G
Δ242-244
K417N
E484K
N501Y
D614G
A701V
Delta
B.1.617.2
1271
T19R
G142D
E156G
Δ157-158
L452R
T478K
D614G
P681R
R682Q
D950N
Omicron
BA.1
1270
A67V
Δ69-70
T95I
G142D
Δ143-145
Δ211
L212I
ins214EPE
G339D
S371L
S373P
S375F
K417N
N440K
G446S
S477N
T478K
E484A
Q493R
G496S
Q498R
N501Y
Y505H
T547K
D614G
H655Y
N679K
P681H
N764K
D796Y
N856K
Q954H
N969K
L981F

**Fig. S6. Characteristics of SARS-CoV-2 S proteins used in the assays based on (a) VSV-SARS-CoV-2 variant pseudoviruses and (b) live authentic SARS-CoV-2.**

The sequence of the Wuhan-Hu-1 isolate SARS-CoV-2 Spike (GenBank: QHD43416.1) was used as reference. Amino acid positions, amino acid descriptions (one letter code) and kind of mutations (substitutions, deletions, insertions) are indicated. NTD, N-terminal domain; RBD, Receptor-binding domain, Δ, deletion; ins, insertion; *, Cytoplasmic domain truncated for the C-terminal 19 amino acids.

**Table S1. BNT162b2 vaccinated individuals analyzed for neutralizing antibody responses**

| Characteristic | BNT162b2$^{2}$ (n=20) | BNT162b2$^{3}$ (n=19) | BNT162b2$^{2}$ + Omi (n=8) | BNT162b2$^{3}$ + Omi (n=10) |
|---|---|---|---|---|
| Sex, n (%) | | | | |
| Male | 10 (50) | 10 (53) | 3 (38) | 7 (70) |
| Female | 10 (50) | 9 (47) | 5 (62) | 3 (30) |
| Age, median (range) | 52 (23-68) | 38 (23-51) | 39 (27-60)° | 32 (23-55)° |
| Age group at vaccination, n (%) | | | | |
| 18-55 yrs | 12 (60) | 19 (100) | 6 (75) | 10 (100) |
| 56-85 yrs | 8 (40) | 0 (0) | 2 (25) | 0 (0) |
| Baseline SARS-CoV-2 status, n (%) | | | | |
| Positive | 0 (0) | 0 (0) | 8 (100)† | 10 (100)† |
| Negative | 20 (100)* | 19 (100)[#] | 0 (0) | 0 (0) |
| Unknown | 0 (0) | 0 (0) | 0 (0) | 0 (0) |
| Interval, median (range) | | | | |
| Days between D1/D2 | 20 (19-21) | ‡ | 40 (21-49) | 38 (20-92) |
| Days until serum draw after D2 | 22 (20-23) | N/A | N/A | n/a |
| Days between D2/D3 | N/A | 202 (181-266) | N/A | 192 (159-255) |
| Days until serum draw after D3 | N/A | 28 (26-30) | N/A | N/A |
| Days between last dose/infection | N/A | N/A | 153 (142-182) | 25 (3-66) |
| Days until serum draw after infection | N/A | N/A | 46 (41-54) | 44 (25-55) |

N/A: not applicable; n/a, not available; D, Dose; Yrs, Years; n, Number.
*, Negative SARS-CoV-2 PCR test at the time of enrollment
[#], No evidence of prior SARS-CoV-2 infection (based on COVID-19 symptoms/signs and SARS-CoV-2 PCR test)
°, Age is estimated based on the indicated year of birth in Table S1
‡, Participants received the primary 2-dose series of BNT162b2 vaccine as part of a governmental vaccination program and the interval between doses was not recorded
†, Omicron infection confirmed at time of recruitment to the research study

**Table S2. BNT162b2 vaccinated individuals analyzed for frequencies of full S protein/RBD-specific $B_{MEM}$ cells**

| Characteristic | BNT162b2$^{2}$ (n=7) | BNT162b2$^{3}$ (n=5) | BNT162b2$^{2}$ + Omi (n=7) | BNT162b2$^{3}$ + Omi (n=5) |
|---|---|---|---|---|
| **Overlap with pVNT/ VNT cohorts (n)** | **n=4** | **n=0** | **n=7** | **n=5** |
| Sex, n (%) | | | | |
| Male | 4 (57) | 4 (80) | 3 (43) | 4 (80) |
| Female | 3 (43) | 1 (20) | 4 (57) | 1 (20) |
| Age, median (range) | 51 (23-80) | 26 (20-69) | 42 (29-60)° | 29 (23-55)° |
| Age group at vaccination, n (%) | | | | |
| 18-55 yrs | 5 (71) | 3 (60) | 6 (86) | 5 (100) |
| 56-85 yrs | 2 (29) | 2 (40) | 1 (14) | 0 (0) |
| Baseline SARS-CoV-2 status, n (%) | | | | |
| Positive | 0 (0) | 0 (0) | 7 (100)† | 5 (100)† |
| Negative | 7 (100)* | 5 (100)# | 0 (0) | 0 (0) |
| Unknown | 0 (0) | 0 (0) | 0 (0) | 0 (0) |
| Interval, median (range) | | | | |
| Days between D1/D2 | 21 (19-23) | 21 (19-23) | 42 (21-49) | 36 (28-42) |
| Days until early blood draw after D2 | 22 (19-23) | N/A | N/A | N/A |
| Days until late blood draw after D2 | 162 (160-167) | N/A | N/A | N/A |
| Days between D2/D3 | N/A | 251 (180-276) | N/A | 189 (159-255) |
| Days until blood draw after D3 | N/A | 84 (81-86) | N/A | N/A |
| Days between last dose/infection | N/A | N/A | 155 (142-182) | 10 (3-27) |
| Days until blood draw after infection | N/A | N/A | 46 (41-54) | 44 (43-47) |

N/A, not applicable; D, Dose; Yrs, Years; n, Number.
*, Negative SARS-CoV-2 PCR test at the time of enrollment
#, No evidence of prior SARS-CoV-2 infection (based on COVID-19 symptoms/signs and SARS-CoV-2 PCR test)
°, Age is estimated based on the indicated year of birth in Table S1
†, Omicron infection confirmed at time of recruitment to the research study

**Table S3. Double and triple BNT162b2-vaccinated individuals with Omicron breakthrough infection**

| Participant ID | YOB | Sex | Vaccination BNT162b2 | Omicron subtype | Dose 1-2 interval | Dose 2-3 interval | Positive test after last vaccination | Blood draw after positive test | Severity (WHO grade) |
|---|---|---|---|---|---|---|---|---|---|
| **1** | 1991 | f | 2 doses | n/a | 42 | N/A | 170 | 46 | 1-2 |
| **2** | 1987 | m | 2 doses | BA.1 | 21 | N/A | 143 | 51 | 1-2 |
| **3** | 1995 | f | 2 doses | BA.1 | 33 | N/A | 151 | 52 | 1-2 |
| **4** | 1977 | m | 2 doses | BA.1 | 42 | N/A | 142 | 43 | 1-2 |
| **5** | 1966 | f | 2 doses | BA.1 | 38 | N/A | 169 | 54 | 1-2 |
| **6** | 1993 | f | 2 doses | n/a | 42 | N/A | 182 | 41 | 1-2 |
| **7** | 1962 | f | 2 doses | n/a | 35 | N/A | 155 | 41 | 1-2 |
| **8** | 1980 | m | 2 doses | n/a | 49 | N/A | 148 | 46 | 1-2 |
| **9** | 1990 | f | 3 doses | n/a | 24 | 243 | 64 | 55 | 1-2 |
| **10** | 1990 | m | 3 doses | n/a | 20 | 233 | 66 | 53 | 1-2 |
| **11** | 1994 | m | 3 doses | n/a | 35 | 213 | 10 | 47 | 1-2 |
| **12** | 1993 | f | 3 doses | BA.1 | 36 | 189 | 3 | 46 | 1-2 |
| **13** | 1999 | m | 3 doses | n/a | 42 | 159 | 27 | 44 | 1-2 |
| **14** | 1991 | m | 3 doses | n/a | 42 | 166 | 20 | 43 | 1-2 |
| **15** | 1969 | m | 3 doses | n/a | 39 | 194 | 22 | 25 | 1-2 |
| **16** | 1972 | f | 3 doses | n/a | 92 | 169 | 35 | 28 | 1-2 |
| **17** | 1972 | m | 3 doses | n/a | 42 | 169 | 44 | 31 | 1-2 |
| **18** | 1962 | m | 3 doses | n/a | 26 | 236 | 112 | 43 | 1-2 |
| | | | | | | **Median** | 65 | 45 | |

YOB, year of birth; m, male; f, female; n/a, not available; N/A, not applicable;

.

**Table S4. $pVN_{50}$ values of sera collected from Omicron-naïve double BNT162b2-vaccinated individuals**

| Clinical trial | Participant ID | $pVN_{50}$ | | | | | | |
|---|---|---|---|---|---|---|---|---|
| | | Wuhan | Alpha | Beta | Delta | Omicron BA.1 | Omicron BA.2 | SARS-CoV-1 |
| BNT162-01 | 19 | 160 | 40 | 10 | 40 | 5 | 5 | 5 |
| | 20 | 320 | n/a | n/a | n/a | 5 | n/a | n/a |
| | 21 | 80 | n/a | n/a | 80 | 5 | n/a | n/a |
| | 22 | 160 | 160 | 40 | 80 | 10 | 20 | 5 |
| | 23 | 320 | 160 | 320 | 320 | 10 | 40 | 5 |
| | 24 | 160 | 80 | 20 | 80 | 5 | 10 | 5 |
| | 25 | 160 | 80 | 20 | 80 | 5 | 10 | 10 |
| | 26 | 320 | 160 | 80 | 160 | 20 | 80 | 20 |
| | 27 | 160 | 80 | 40 | 40 | 5 | 40 | 5 |
| | 28 | 160 | 80 | 20 | 80 | 10 | 20 | 5 |
| | 29 | 160 | 40 | 10 | 80 | 5 | 5 | 5 |
| | 30 | 160 | 80 | 10 | 40 | 5 | 10 | 5 |
| | 31 | 160 | 80 | 160 | 80 | 5 | 20 | 5 |
| | 32 | 80 | 40 | 10 | 40 | 5 | 5 | 5 |
| | 33 | 160 | 160 | 20 | 80 | 5 | 20 | 5 |
| | 34 | 640 | 320 | 40 | 160 | 10 | 20 | 5 |
| | 35 | 160 | 320 | 80 | 160 | 10 | 20 | 5 |
| | 36 | 160 | 80 | 20 | 80 | 5 | 20 | 5 |
| | 37 | 20 | 20 | 5 | 5 | 5 | 5 | 5 |
| | 38 | 160 | 80 | 10 | 80 | 5 | 5 | 5 |

n/a, not available due to lack of serum.

**Table S5. pVN50 values of sera collected from Omicron-naïve triple BNT162b2-vaccinated individuals**

| Clinical trial | Participant ID | pVN$_{50}$ | | | | | | |
|---|---|---|---|---|---|---|---|---|
| | | Wuhan | Alpha | Beta | Delta | Omicron BA.1 | Omicron BA.2 | SARS-CoV-1 |
| BNT162-17 | 39 | 160 | 320 | 160 | 160 | 80 | 160 | 5 |
| | 40 | 320 | 640 | 320 | 640 | 320 | 320 | 10 |
| | 41 | 640 | 640 | 320 | 320 | 160 | 320 | 40 |
| | 42 | 5120 | 5120 | 1280 | 2560 | 1280 | 1280 | 40 |
| | 43 | 320 | 640 | 160 | 320 | 160 | 160 | 20 |
| | 44 | 640 | 640 | 80 | 640 | 320 | 160 | 20 |
| | 45 | 320 | 640 | 160 | 320 | 160 | 160 | 10 |
| | 46 | 320 | 640 | 320 | 320 | 160 | 160 | 10 |
| | 47 | 320 | 640 | 320 | 320 | 160 | 160 | 20 |
| | 48 | 160 | 320 | 80 | 160 | 40 | 80 | 20 |
| | 49 | 320 | 1280 | 160 | 320 | 160 | 60 | 20 |
| | 50 | 1280 | 5120 | 640 | 1280 | 640 | 640 | 80 |
| | 51 | 40 | 40 | 20 | 20 | 5 | 40 | 5 |
| | 52 | 320 | 640 | 320 | 320 | 80 | 160 | 20 |
| | 53 | 160 | 320 | 160 | 320 | 80 | 160 | 20 |
| | 54 | 320 | 640 | 320 | 320 | 320 | 320 | 20 |
| | 55 | 640 | 640 | 320 | 320 | 160 | 320 | 40 |
| | 56 | 2560 | 5120 | 640 | 1280 | 640 | 640 | 80 |
| | 57 | 320 | 640 | 160 | 640 | 160 | 320 | 20 |

**Table S6. pVN$_{50}$ values of sera collected from double and triple BNT162b2-vaccinated individuals after confirmed Omicron breakthrough infection**

| Participant ID | pVN$_{50}$ | | | | | | |
|---|---|---|---|---|---|---|---|
| | Wuhan | Alpha | Beta | Delta | Omicron BA.1 | Omicron BA.2 | SARS-CoV-1 |
| 1* | 960 | 1920 | 960 | 960 | 960 | 480 | 120 |
| 2* | 960 | 1920 | 960 | 960 | 1920 | 480 | 120 |
| 3* | 960 | 1920 | 960 | 480 | 960 | 960 | 120 |
| 4* | 480 | 480 | 480 | 240 | 480 | 480 | 120 |
| 5* | 240 | 480 | 240 | 120 | 480 | 120 | 5 |
| 6* | 1920 | 1920 | 1920 | 1920 | 960 | 960 | 40 |
| 7* | 960 | 1920 | 960 | 960 | 960 | 960 | 40 |
| 8* | 480 | 960 | 480 | 480 | 960 | 480 | 20 |
| 9# | 1920 | 3840 | 1920 | 1920 | 1920 | 960 | 120 |
| 10# | 960 | 1920 | 960 | 480 | 480 | 480 | 120 |
| 11# | 3840 | 3840 | 3840 | 3840 | 1920 | 1920 | 960 |
| 12# | 960 | 1920 | 960 | 960 | 960 | 960 | 120 |
| 13# | 480 | 1920 | 960 | 480 | 480 | 480 | 20 |
| 14# | 1920 | 1920 | 960 | 960 | 960 | 1920 | 40 |
| 15# | 960 | 1920 | 960 | 960 | 1920 | 480 | 80 |
| 16# | 960 | 1920 | 480 | 960 | 480 | 480 | 5 |
| 17# | 480 | 960 | 480 | 480 | 480 | 480 | 60 |
| 18# | 1920 | 3840 | 3840 | 3840 | 3840 | 1920 | 120 |

*, participant received two doses of BNT162b2 prior to Omicron infection
#, participant received three doses of BNT162b2 prior to Omicron infection

**Table S7. $VN_{50}$ values of sera collected from Omicron-naïve double BNT162b2-vaccinated individuals**

| Clinical trial | Participant ID | $VN_{50}$ | | | |
|---|---|---|---|---|---|
| | | Wuhan | Beta | Delta | Omicron |
| BNT162-01 | 19 | 226 | 5 | 20 | 5 |
| | 20 | 320 | 14 | 80 | 5 |
| | 21 | 226 | 5 | 40 | 5 |
| | 22 | 905 | 28 | 80 | 7 |
| | 23 | 640 | 113 | 226 | 10 |
| | 24 | 320 | 14 | 20 | 5 |
| | 25 | 453 | 28 | 80 | 14 |
| | 26 | 1810 | 80 | 160 | 28 |
| | 27 | 453 | 28 | 40 | 7 |
| | 28 | 320 | 14 | 80 | 7 |
| | 29 | 226 | 5 | 20 | 5 |
| | 30 | 453 | 14 | 28 | 5 |
| | 31 | 320 | 28 | 40 | 5 |
| | 32 | 160 | 14 | 14 | 5 |
| | 33 | 640 | 20 | 80 | 5 |
| | 34 | 320 | 20 | 20 | 7 |
| | 35 | 320 | 28 | 40 | 7 |
| | 36 | 640 | 20 | 56 | 5 |
| | 37 | 40 | 5 | 5 | 5 |
| | 38 | 320 | 7 | 40 | 5 |

**Table S8. $VN_{50}$ titer values of sera collected from Omicron-naïve triple BNT162b2-vaccinated individuals**

| Clinical trial | Participant ID | $VN_{50}$ titer | | | |
|---|---|---|---|---|---|
| | | Wuhan | Beta | Delta | Omicron BA.1 |
| BNT162-17 | 39 | 226 | 80 | 226 | 40 |
| | 40 | 905 | 453 | 320 | 226 |
| | 41 | 640 | 226 | 226 | 160 |
| | 42 | 3620 | 1280 | 1810 | 453 |
| | 43 | 226 | 226 | 160 | 40 |
| | 44 | 226 | 80 | 160 | 57 |
| | 45 | 640 | 113 | 226 | 80 |
| | 46 | 453 | 226 | 453 | 80 |
| | 47 | 640 | 226 | 226 | 160 |
| | 48 | 160 | 80 | 113 | 28 |
| | 49 | 640 | 80 | 320 | 80 |
| | 50 | 3620 | 905 | 1280 | 453 |
| | 51 | 28 | 10 | 10 | 5 |
| | 52 | 453 | 113 | 226 | 40 |
| | 53 | 226 | 113 | 226 | 57 |
| | 54 | 905 | 160 | 453 | 160 |
| | 55 | 1280 | 160 | 226 | 113 |
| | 56 | 3620 | 905 | 1810 | 1280 |
| | 57 | 905 | 226 | 320 | 80 |

**Table S9. $VN_{50}$ titer values of sera collected from double and triple BNT162b2-vaccinated individuals after confirmed Omicron breakthrough infection**

| Participant ID | $VN_{50}$ titer | | | |
|---|---|---|---|---|
| | Wuhan | Beta | Delta | Omicron BA.1 |
| 1* | 453 | 453 | 453 | 640 |
| 2* | 640 | 905 | 1280 | 640 |
| 3* | 453 | 640 | 320 | 640 |
| 4* | 160 | 453 | 226 | 320 |
| 5* | 80 | 113 | 160 | 453 |
| 6* | 1280 | 905 | 1280 | 640 |
| 7* | 905 | 905 | 640 | 453 |
| 8* | 226 | 320 | 320 | 320 |
| 9# | 453 | 1280 | 1280 | 640 |
| 10# | 453 | 640 | 640 | 453 |
| 11# | 1810 | 1810 | 2560 | 1810 |
| 12# | 453 | 905 | 1280 | 453 |
| 13# | 640 | 640 | 320 | 453 |
| 14# | 640 | 640 | 905 | 640 |
| 15# | 320 | 640 | 453 | 453 |
| 16# | 905 | 226 | 453 | 226 |

*, participant received two doses of BNT162b2 prior to Omicron infection
#, participant received three doses of BNT162b2 prior to Omicron infection

**Table S10. Individuals vaccinated with other approved COVID-19 vaccines or mixed regimens after subsequent Omicron breakthrough infection**

| Participant ID | YOB | Sex | Vaccination | Omicron subtype | Dose 1-2 interval | Dose 2-3 interval | Positive test after last vaccination | Blood draw after positive test | Severity (WHO grade) |
|---|---|---|---|---|---|---|---|---|---|
| **58** | 1960 | f | AZ/BNT | n/a | 62 | N/A | 142 | 46 | 1-2 |
| **59** | 1955 | m | AZ/BNT | n/a | 68 | N/A | 135 | 45 | 1-2 |
| **60** | 1983 | m | J&J | n/a | N/A | N/A | 161 | 44 | 1-2 |
| **61** | 1962 | m | MOD$^3$ | BA.1 | 42 | 172 | 3 | 35 | 1-2 |
| **62** | 1993 | f | MOD$^2$ | n/a | 42 | N/A | 169 | 44 | 1-2 |
| **63** | 1981 | m | J&J/BNT | n/a | 138 | N/A | 45 | 40 | 1-2 |
| **64** | 1989 | m | AZ/BNT/MOD | BA.1 | 68 | 154 | 9 | 43 | 1-2 |
| **65** | 1994 | m | MOD$^2$/BNT | n/a | 28 | 252 | 22 | 40 | 1-2 |
| **66** | 1990 | m | MOD$^2$/BNT | n/a | 42 | 154 | 13 | 42 | 1-2 |
| **67** | 1972 | m | MOD$^2$/BNT | n/a | 28 | 256 | 45 | 31 | 1-2 |
| | | | | | | **Median:** | 45 | 43 | |

OB, year of birth; m, male; f, female; n/a, not available; N/A, not applicable;
AZ, AstraZeneca AZD1222; BNT, BioNTech/Pfizer BNT162b2; J&J, Johnson & Johnson Ad26.COV2.S; MOD, Moderna mRNA-1273; BNT$^4$, BNT162b2 four-dose series; MOD$^2$, mRNA-1273 two-dose series; MOD$^3$, mRNA-1273 three-dose series
.

**Table S11. $pVN_{50}$ titer values of sera collected from individuals vaccinated with other approved COVID-19 vaccines or mixed regimens after subsequent Omicron breakthrough infection**

| Participant ID | $pVN_{50}$ titer | | | | | | |
|---|---|---|---|---|---|---|---|
| | Wuhan | Alpha | Beta | Delta | Omicron BA.1 | Omicron BA.2 | SARS-CoV-1 |
| 58 | 1920 | 3840 | 1920 | 960 | 960 | 960 | 240 |
| 59 | 1920 | 3840 | 960 | 1920 | 1920 | 960 | 10 |
| 60 | 120 | 480 | 120 | 240 | 120 | 60 | 60 |
| 61 | 7680 | 15360 | 7680 | 3840 | 15360 | 3840 | 480 |
| 62 | 120 | 120 | 30 | 30 | 60 | 60 | 5 |
| 63 | 480 | 1920 | 960 | 960 | 480 | 480 | 120 |
| 64 | 480 | 1920 | 960 | 480 | 240 | 480 | 60 |
| 65 | 480 | 240 | 60 | 240 | 60 | 120 | 30 |
| 66 | 1920 | 1920 | 960 | 1920 | 960 | 960 | 60 |
| 67 | 3840 | 15360 | 7680 | 7680 | 3840 | 3840 | 120 |

**Table S12. Frequencies of S protein/RBD-specific $B_{MEM}$ cells for all individuals analyzed**

| Study | Participant ID | Wuhan Spike$^+$ $B_{MEM}$s | Alpha Spike$^+$ $B_{MEM}$s | Delta Spike$^+$ $B_{MEM}$s | Omicron Spike$^+$ $B_{MEM}$s | Wuhan RBD$^+$ $B_{MEM}$s | Alpha RBD$^+$ $B_{MEM}$s | Delta RBD$^+$ $B_{MEM}$s | Omicron RBD$^+$ $B_{MEM}$s |
|---|---|---|---|---|---|---|---|---|---|
| Investigator Study | 1* | 3.87 | 3.16 | 3.16 | 3.79 | 0.49 | 0.52 | 0.50 | 0.47 |
| | 2* | 1.09 | 0.88 | 0.94 | 0.98 | 0.62 | 0.66 | 0.46 | 0.47 |
| | 4* | 0.94 | 0.81 | 0.72 | 1.00 | 0.23 | 0.33 | 0.20 | 0.21 |
| | 5* | 0.71 | 0.54 | 0.63 | 0.78 | 1.96 | 1.96 | 1.38 | 1.49 |
| | 6* | 2.15 | 1.83 | 1.91 | 2.09 | 1.15 | 1.17 | 1.10 | 0.89 |
| | 7* | 2.57 | 2.09 | 2.25 | 2.53 | 2.06 | 2.13 | 1.86 | 1.59 |
| | 8* | 0.45 | 0.33 | 0.34 | 0.62 | 0.21 | 0.26 | 0.21 | 0.21 |
| | 11# | 0.73 | 0.73 | 0.93 | 0.90 | 0.37 | 0.29 | 0.23 | 0.34 |
| | 12# | 1.38 | 1.34 | 1.86 | 1.87 | 1.12 | 0.93 | 0.72 | 1.06 |
| | 13# | 1.67 | 1.65 | 2.15 | 2.05 | 1.12 | 0.94 | 0.68 | 1.06 |
| | 14# | 1.35 | 1.33 | 1.68 | 1.62 | 0.94 | 0.75 | 0.72 | 0.91 |
| | 15# | 2.75 | 2.77 | 3.40 | 3.11 | 2.53 | 2.22 | 1.68 | 2.41 |
| | 16# | 0.58 | 0.61 | 0.82 | 0.73 | 0.50 | 0.44 | 0.29 | 0.42 |
| BNT162-01 Early | 35 | 0.28 | 0.18 | 0.19 | 0.27 | 0.11 | 0.12 | 0.13 | 0.05 |
| | 38 | 0.33 | 0.18 | 0.25 | 0.29 | 0.08 | 0.09 | 0.09 | 0.03 |
| | 41 | 0.19 | 0.13 | 0.14 | 0.18 | 0.05 | 0.08 | 0.07 | 0.04 |
| | 47 | 0.31 | 0.20 | 0.16 | 0.31 | 0.07 | 0.10 | 0.08 | 0.02 |
| | 69 | 0.19 | 0.16 | 0.16 | 0.18 | 0.07 | 0.11 | 0.08 | 0.04 |
| | 70 | 0.59 | 0.39 | 0.45 | 0.44 | 0.23 | 0.26 | 0.19 | 0.15 |
| | 71 | 0.52 | 0.37 | 0.46 | 0.46 | 0.16 | 0.23 | 0.17 | 0.07 |
| BNT162-01 Late | 38 | 0.86 | 0.87 | 1.17 | 1.27 | 0.46 | 0.35 | 0.23 | 0.39 |
| | 41 | 0.37 | 0.39 | 0.67 | 0.60 | 0.23 | 0.18 | 0.12 | 0.18 |
| | 68 | 0.31 | 0.36 | 0.55 | 0.47 | 0.18 | 0.13 | 0.06 | 0.09 |
| | 69 | 1.97 | 2.05 | 2.67 | 2.65 | 0.68 | 0.55 | 0.44 | 0.66 |
| | 70 | 1.14 | 1.27 | 1.71 | 1.64 | 0.43 | 0.28 | 0.17 | 0.31 |
| BNT162-14 | 71 | 0.96 | 0.75 | 0.75 | 0.89 | 0.30 | 0.35 | 0.27 | 0.15 |
| | 72 | 0.62 | 0.44 | 0.45 | 0.62 | 0.23 | 0.26 | 0.28 | 0.16 |
| | 73 | 1.92 | 1.35 | 1.55 | 1.48 | 0.62 | 0.62 | 0.59 | 0.50 |
| | 74 | 1.36 | 1.06 | 0.94 | 1.25 | 0.41 | 0.47 | 0.40 | 0.25 |
| | 75 | 2.01 | 1.58 | 1.58 | 1.86 | 0.64 | 0.68 | 0.62 | 0.49 |

*, subject received two doses of BNT162b2 prior to Omicron infection

#,subject received three doses of BNT162b3 prior to Omicron infection
